# Local consequences of FMRP loss: cell type-specific mRNA and protein profiles of *Fmr1*-KO synapses

**DOI:** 10.64898/2026.11.11.765892

**Authors:** Kristina Desch, Selma Aghabashlou Saisan, Susanne tom Dieck, Nicole Fürst, Belquis Nassim-Assir, Julian D. Langer, Marcel Jüngling, Eva Kaulich, Erin M. Schuman

## Abstract

Brain function relies on billions of neurons interconnected by synapses with unique molecular composition specialized for cell type, brain region, and their role in circuit dynamics. Pathological synaptic dysfunctions – so-called synaptopathies – underlie many neurodevelopmental disorders (NDDs), including Fragile X Syndrome (FXS). While synaptic deficits are well documented in FXS, molecular studies have largely relied on bulk brain homogenates or synaptic fractions, which typically represent “average” molecular identities, potentially masking synapse type-specific alterations. To provide synapse type resolution, we use fluorescence-activated synaptosome sorting coupled to RNA-sequencing and mass spectrometry to isolate and profile cortical excitatory and inhibitory synapses of hemizygous and heterozygous *Fmr1*-deficient mice. We found that full or partial FMRP loss leads to cell type- and compartment-specific molecular remodeling (syndive.org), affecting mRNAs and proteins related to synaptic organization and metabolic pathways. Strikingly, only a minority of dysregulated mRNAs/proteins were shared across excitatory and inhibitory synapses - the substantial majority were uniquely regulated at one subtype. Indeed, FMRP loss often resulted in opposing protein changes in each synapse type. Changes in mRNA or protein abundance were also not well coordinated, consistent with substantial post-transcriptional regulation, including changes to the translational and ubiquitin-dependent protein degradation machinery. For example, our data suggest a disturbed, cell type-distinct coordination of protein degradation via CAND1 as one key consequence of FMRP loss. Furthermore, we observed differences in male (*Fmr1*^-/y^) and female (*Fmr1*^-/+^) transcriptomes, where, for example, FMRP dosage was coupled to reciprocal regulation of the NDD-risk gene *Trpc4* in both synapse types.

## Introduction

The mammalian brain contains billions of neurons that communicate via highly specialized synaptic connections. Synapses differ across brain regions and neuronal cell types, displaying distinct complements of mRNAs [1, 2] and proteins [3, 4] that support their functional specialization. Excitatory and inhibitory synapses constitute two major functional classes that promote or constrain neuronal activity, respectively, and together maintain balanced circuit functions. This molecular diversity enables spatially-controlled regulation of synaptic transmission and plasticity [5] and thereby contributes to higher-order functions, including learning, memory, and perception [6]. Conversely, alterations in these molecular signatures can impair information processing and activity-dependent remodeling [7, 8]. Synaptic dysfunction is therefore indeed a common feature of many neurodevelopmental and neurodegenerative disorders, frequently conceptualized as synaptopathies [9].

Fragile X syndrome (FXS) is a major monogenic cause of intellectual disability (ID) and often presents with autism spectrum disorder (ASD) [9]. It is typically caused by expansion of a CGG trinucleotide repeat within the 5′ untranslated region of the fragile X messenger ribonucleoprotein 1 (*FMR1*) gene, resulting in transcriptional silencing and consequent loss of the encoded FMR protein (FMRP; [10, 11]. FMRP regulates the transport, stability, and translation of neuronal mRNAs and has been proposed to control ribosome stalling on selected transcripts [12–17]. Beyond its role as RNA-binding protein, FMRP has also been reported to alter ion channel properties via direct interaction [18]. Through all these functions, FMRP is well poised to contribute to spatio-temporal regulation of protein synthesis in neuronal processes and at synapses.

Loss of FMRP causes diverse synaptic dysfunctions, including cell type- and circuit-dependent alterations in excitation-inhibition balance [19, 20], an increased prevalence of thin and immature dendritic spines [21–24], and impaired synaptic plasticity [25, 26, 12, 27], particularly enhanced metabotropic glutamate receptor-dependent long-term depression in the hippocampus [25] and reduced long-term potentiation in cortical circuits of *Fmr1*-deficient mice [26]. Clinically, humans with FXS commonly present with ID, altered sensory processing, hyperactivity, anxiety, and autistic features [28]. Disease phenotypes are generally more severe in males, who typically lack FMRP, whereas heterozygous females show variability in reduced FMRP expression and phenotypic severity, most likely owing to X-chromosome inactivation [28, 29]. Further, environmental influences in combination with the individual genetic susceptibility shape disease expression and contribute to phenotypic heterogeneity [30, 31].

Studies using a functional endpoint mouse model, the *Fmr1*-knockout (*Fmr1*-KO), have implicated dysregulated mRNA transport [13], translation [15], and protein homeostasis in FXS pathogenesis [32, 27, 33]. The consequences of FMRP loss, however, present in a highly context-dependent manner and vary across brain regions, developmental stages, sex, and experimental paradigm [34, 27, 35]. The heterogeneity observed might reflect a role of FMRP in fine-tuning the balance between protein synthesis and degradation rather than acting as a uniform translational switch, a concept previously introduced for FXS and other ASD-/ID-mutations (for review: [36, 37]).

It is important to note that bulk samples combine molecules from multiple neuronal and non-neuronal cell types, obscuring differences between somatic, dendritic, axonal, and synaptic compartments [15, 34, 32, 27, 33]. As such, most previous studies have faced an „averaging problem“, preventing molecular alterations from being assigned to defined neuronal populations or subcellular compartments. These limitations are particularly relevant in FXS, where opposing or cell type-specific responses can contribute to excitation-inhibition imbalance [19, 20] and are most likely masked in bulk measurements. Single-cell and single-nucleus transcriptomic approaches [8, 38] or cell type-specific RiboTag-Seq [39] provide cellular resolution but may primarily capture somatic RNA and cannot resolve the molecular composition of distinct synapse subtypes.

Here, following fluorescence-activated synaptosome sorting (FASS) coupled to RNA-sequencing (RNA-Seq) and data-independent acquisition mass spectrometry (DIA-MS), we analyzed purified synaptosomes of cortical excitatory and inhibitory neurons from young adult (7 weeks) male hemizygous *Fmr1*^−/y^ and female heterozygous *Fmr1*^−/+^ mice. This approach enables us to identify molecular alterations that are shared across synapse types and sexes, distinguish them from synapse type-specific effects, and determine to what extent changes of the local proteome correspond to changes in synaptic mRNA abundance. In-depth multi-omics analyses revealed population-specific regulatory patterns in both the transcriptome and proteome. Moreover, we found that these mechanisms operate at different targets, only a minority of which have previously been identified as FMRP-bound transcripts, suggesting that FMRP loss engages distinct regulatory mechanisms across synaptic populations and omics-paradigms.

## Methods

### Experimental model and animal details

All procedures involving animal treatment and care were conducted in conformity with the institutional guidelines that are in compliance with national and international laws and policies (DIRECTIVE 2010/63/EU; German animal welfare law; FELASA guidelines). The animals were euthanized according to annex 2 of § 2 Abs. 2 Tierschutz-Versuchstier-Verordnung and animal numbers were reported to the local authorities (Regierungspräsidium Darmstadt). Mice were housed under a 12 hour light/dark cycle at 20-24 °C ambient temperature and 55 ± 10% relative humidity. Food and water were provided *ad libidum*.

### Breeding strategy and animals used for transcriptomics and proteomics

To profile the synaptic transcriptome and proteome of inhibitory and excitatory neurons in the FXS model, we crossed homozygous Gad2-(Gad2^tm2(cre)Zjh^/J; IMSR_JAX:010802; [40]) or Nex-Cre (Nave lab, MPI for multidisciplinary sciences, Göttingen; [41]) driver line females with floxed Synaptophysin-tdTomato reporter line (B6;129S-Gt(ROSA)26So^rtm34.1(CAG-Syp/tdTomato)Hze^/J; IMSR_JAX:012570, [42]) males to obtain Nex-Cre::SypTOM^+^ or Gad2-Cre::SypTOM^+^ mice to label presynaptic terminals. The offspring, heterozygous in both loci, was further crossed by sibling matings to generate double homozygous Nex-Cre::SypTOM^+^ animals and Gad2-Cre::SypTOM^+^ animals homozygous for at least one of the alleles. X-Cre::SypTOM^+^ males were then crossed with *Fmr1*^-/+^ females (B6.129P2-*Fmr1*^tm1Cgr^/J; IMSR_JAX:003025; [43]) to generate littermate offspring with the respective synapse populations labeled in a *Fmr1*-KO and *Fmr1*-WT animal. After some cases of reporter germline activation in the Gad2-Cre::SypTOM^+^ animals, we switched to crossing the Gad2-Cre line with *Fmr1*-KO animals first to obtain *Fmr1*^-/+^::Gad2-Cre^ki/ki^ females that were crossed to homozygous SypTOM males to obtain the experimental animals as littermates. In all cases, the *Fmr1* alleles determining KO or WT classification were derived from the female and the SypTOM allele was inherited from the male parent. All experiments were performed on adult mice of 7 weeks. Animals of different sex and hence genotype (male study: wildtype WT, *Fmr1*^+/y^ and hemizygous KO, *Fmr1*^-/y^; female study: wildtype WT *Fmr1*^+/+^ and heterozygous KO, *Fmr1*^-/+^; all animals were heterozygous for the X-Cre and SypTOM alleles) were used in the separate omics analyses.

### Brain dissection, cortex tissue and synaptosome preparation

All mice were sacrificed by decapitation under deep isoflurane anesthesia, their brains dissected and placed on an iced Petri dish. One brain hemisphere was dissected for PFA-fixation. Fixation was performed for 2 hours at RT using 4% PFA, 4% sucrose in phosphate-buffered saline (PBS; Gibco, Thermo Fisher Scientific, Waltham, MA, USA; #cat 10010023) at pH 7.4. Followed by washes with PBS, first 15% sucrose in PBS was applied at 4°C overnight. The next day, brains were transferred into 30% sucrose in PBS at 4°C overnight and kept at -80°C until use. For synaptosome preparation (according to [44] with modifications of [3]), the second cortex hemisphere was collected in 3 ml ice-cold GM buffer (0.25 M sucrose, 5 mM tris hydroxymethyl aminomethane [TRIS; Sigma-Aldrich, St. Louis, MO, USA; cat# 252859]-HCl pH 7.5, 0.1 mM EDTA supplemented with 1:80 RNAse (RI; SUPERase-In RNAse Inhibitor, Invitrogen, cat# AM2696) and 1:750 protease inhibitor (PR; Protease Inhibitor CocktailSet III, EMD, Merck Millipore, Burlington, MA, USA; cat# 539134-10ml) and transferred to a glass Dounce Tissue Grinder (WHEATON®, 7 ml) for homogenization using 10 strokes of the “loose” and 10 of the “tight” pestle. The homogenate was then centrifuged for 10 minutes at 4°C at 1,000 x g in a swinging bucket rotor. The supernatant (S1) was layered onto a density gradient (23%, 10%, and 3% Percoll [Cytiva, Merck Millipore, Burlington, MA, USA; cat# GE17-0891-01] in GM), which was centrifuged in an Avanti J-26S XPI centrifuge with a JA-25.50 rotor (Beckman Coulter, Brea, CA, USA) for 5 minutes at 4°C at 32,500 x g with maximum acceleration and minimum deceleration. The F2/3 fraction, also referred to as unsorted (crude) synaptosomes, was collected with a 5ml syringe and a long, blunt cannula (∼1,200 µl at the interface of 23% and 10% Percoll). Before it was subjected to FASS, it was diluted at least in a 1:25 ratio with GM buffer (with RI and PI). For bulk proteomics analyses, aliquots of the original cortex homogenate and F2/3 fraction were snap-frozen and kept at -80°C until MS sample preparation.

### Fluorescence-activated synaptosome sorting (FASS)

FASS was performed on a FACS Aria Fusion (BD Biosciences, San Jose, CA, USA) running BD FACSDiva^TM^ (BD Biosciences, San Jose, CA, USA), as previously described [3, 1, 4, 2]. In brief, the following system settings were used: doublet particle exclusion via SSC-H and SSC-W, 488 nm laser (FM4-64), 561 nm laser (tdTomato signal), sort precision (0-16-0), FSC (317 V), SSC (488/10 nm, 370 V), PE “tdTomato” (586/15 nm, 470 V), PerCP “FM4-64” (695/40 nm, 470 V), thresholds (FSC = 200, FM4-64 = 700). Samples were analyzed and sorted at ∼10,000-15,000 events/s and a flow rate below 3. BD FACSFlow (BD Biosciences, San Jose, CA, USA; cat# 12756528) was used as a sorting buffer and synaptosomes were sorted using a 70 µm nozzle. Collection was performed by filtration onto a Whatman GF/F glass microfiber filter (Cytiva, Marlborough, MA, USA; cat# 1825-090) supported by a Whatman PE drain-disk (Merck Millipore, Burlington, MA, USA; cat# WHA231100). Using a Miltex biopsy punch (4 mm; Integra LifeSciences, York, PA, USA; cat# 15110-40-X), they were punched out on top of one another and prior sorting, they were washed with 700 µl PBS containing RNAsecure (1:10; Thermo Fisher Scientific, Waltham, MA, USA; cat# AM7006) and Protease inhibitor cocktail III (Millipore, Burlington, MA, USA 1:750; cat# 539134). For each analysis, 2 million synaptosomes were collected on filter, which was washed with 1 ml RNAse-free PBS for transcriptomics or 50 mM triethylammonium bicarbonate (TEAB; Sigma-Aldrich, St. Louis, MO, USA; cat# 18597-100ML) for proteomics. Purified synaptosome (referred to as P3) fractions were sorted based on the presence of a membrane dye (FM4-64; Thermo Fisher Scientific, Waltham, MA, USA; cat# T13320) and tdTomato signal, while control (referred to as notP3) fractions were the remaining particles that did not match the P3 gate in the sorting layout after doublet particle exclusion. After sorting, filters containing the purified synaptosomes or control particles were stored at -80 °C until sample preparation. Quantification was done using the BD FACSDiva^TM^ software (BD Biosciences, San Jose, CA, USA). Visualization of FACS contour plots was performed in FlowJo^TM^ (v10.10.1; BD Biosciences, San Jose, CA, USA).

### Library preparation and sequencing of sorted synaptosomes

Sequencing libraries were prepared with n = 5 animals per condition (n*_Fmr1-/y, Fmr1+/y, Fmr1-/+, Fmr1+/+_* = 5 for Nex^+^ and Gad2^+^ and P3 and notP3 each). Sequencing libraries of sorted synaptosomes were prepared using a previously established low-input sample protocol modified from a subcellular RNA sequencing protocol [45, 1, 2]. For membrane lysis and DNA digestion, synaptosomes were resuspended from the sorting filter by adding 7.5 µl of 1 mM dithiothreitol (DTT; PanReac AppliChem, Darmstadt, Germany; cat# A3668), 1U/µl Recombinant RNAse Inhibitor (Takara Bio, Kusatsu, Shiga, Japan; cat# 2313A) and 1x SingleShot Lysis Buffer with 1x DNAse solution (BioRAD, Hercules, CA, USA; cat# 1725080) to each sample and incubated for 20 min at 25°C, followed by 5 min at 75°C for enzyme inactivation. Next, RT_index primers carrying a sample-specific index and UMI sequences (see **Table S23**, final concentration 1 mM) were hybridized to polyA stretches by adding 4.6 µl of DTT (10 mM), dNTPs (1 mM; ThermoFisher Scientific, Waltham, MA, USA; R1121), dCTPs (1.5 mM; ThermoFisher Scientific, Waltham, MA, USA; cat# R0151) and Recombinant RNAse Inhibitor (1 U/µl) to each sample. The samples were incubated at 65°C for 5 min and afterwards immediately kept on ice for at least 1 min. Reverse transcription and template switching were carried out by adding 10.9 µl of the reverse transcriptase SuperScript IV (10 U/µl) with SuperScript RT buffer (1x; ThermoFisher Scientific, Waltham, MA, USA; cat# 18090050), MgCl_2_ (6 mM) (Invitrogen, Carlsbad, CA, USA; cat# AM9530G), Betaine (1 M, Merck, Burlington, MA, USA; cat# B0300-5VL), and template switch oligo (1 µM, IDT, **Table S23**) and incubating the sample for 10 min at 55 °C, followed by enzyme inactivation for 10 min at 80 °C. For PCR, 23 µl of the KAPA HiFi HotStart Ready Mix (1x) (Roche, Mannheim, Germany; cat# 07958935001) and PCR primer (0.1 µM, IDT, **Table S23**) were added to each sample. Samples were incubated for 3 min at 98°C for activating the DNA polymerase and went through 18 cycles of incubation at 98°C for 20 s for DNA denaturation, 67°C for 15 s for annealing, and 72°C for 6 min of extension. Lastly, the samples were incubated at 72°C for 5 min for an additional extension phase and kept on ice. After PCR, the sorting filter was removed by transferring the sample to a centrifuge tube filter (Spin-X Centrifuge 764 Tube Filter 0.45 µM Pore CA Membrane in 2.0 ml Tube; Corning, Corning, NY, USA, cat# 8162), centrifuging for 1 min at 3000 g, and transferring the volume to 0.2 ml PCR tubes. For DNA purification and selecting large fragments, 28.8 µl (0.6x of the sample volume) of AMPure XP beads (Beckman Coulter, Brea, CA, USA; cat# A63881) were added to each sample following manufacturer’s instructions. Next, libraries with different RT primer indexes were pooled and indexed using the Nextera XT DNA library prep kit (Illumina, San Diego, CA, USA) with a custom P5 primer (IDT, **Table S23**). Paired-end sequencing was performed using the NextSeqTM 2000 system (Illumina, San Diego, CA, USA) with NextSeqTM 1000/2000 P3 cartridges (100 cycles) and a custom Read 1 sequencing primer (IDT, **Table S23**), using the following read parameters: Read 1 = 26 bp, Read2 = 82 bp, Read 1 Index = 8 bp, Read 2 Index = 0.

### Preprocessing methods for RNA-Seq of synaptosome datasets

Barcoded synaptosome RNA-Seq data was processed using a previously published custom bash pipeline [1, 4, 2]. Raw BCL files were demultiplexed with ‘bcl2fastq’ (https://github.com/s-andrews/fastqc) generating paired-end FASTQ files. Paired-end reads were aligned to the *Mus musculus* mm39 reference genome and annotation from the UCSC Genome Browser using STAR (v2.7.11b; [46]) in STARsolo mode with ‘--soloType CB_UMI_Simple’. Barcode error correction was performed allowing one mismatch against a predefined whitelist (‘--soloCBmatchWLtype 1MM’) and UMIs were deduplicated with a one-mismatch scheme across all genes (‘--soloUMIdedup 1MM_All’). Gene expression quantification was performed with ‘--soloFeatures GeneFull_Ex50pAS’. Resulting BAM files, UMI count matrices, and alignment logs were organized for downstream analyses.

### Differential expression analysis of the synaptic transcriptome

Differential expression analysis was conducted in R 4.5.2 using DESeq2 (v1.50.2; [47]) with protein-coding RNAs based on the *Mus musculus* mm39 reference genome (NCBI RefSeq assembly: GCF_000001635.27). Transcripts were filtered with mean count > 15 across samples to remove potential noise. Counts originating from mitochondrially encoded genes and high-confidence microglial contaminants [48, 1] were further removed to focus on localized neuronal transcripts expressed in the cell nucleus. Nextera PCR batch effects were corrected using COMBATseq (sva package v3.58.0; [49]). Downstream differential expression analysis for synaptic purity, i.e. P3 vs. notP3, was conducted with *Fmr1^+/+^* and *Fmr1^+/y^* samples for each synapse type individually using DESeq2. A DESeqDataSet object using the factor “Gate” – encoding the P3 and notP3 samples – was set for the design formula (design = ∼ Gate). Differential expression analysis for determining dysregulated transcripts between *Fmr1^-/y^* vs. *Fmr1^+/y^*, *Fmr1^-/+^*vs. *Fmr1^+/+^* and *Fmr1^+/+^* vs. *Fmr1^+/y^* was conducted for each synapse type separately. A DESeqDataSet object with the factor “Idshort” – encoding the individual *Fmr1* genotypes – was set as the design formula (design = ∼ Idshort). Differential expression analysis was performed using a negative binomial generalized linear model with the default Wald test and local dispersion fit. Log_2_ fold-changes were shrunk using the apeglm method [50]. DESeq2 p-values were adjusted using independent hypothesis weighting (IHW package v1.38.0) at a significance level of 0.1 [51]. The weight of each test was modeled using the mean of normalized counts as a covariate. Transcripts were defined as significantly dysregulated using padj (FDR) < 0.05. For principal component analysis (PCA) of synaptic transcriptomes, counts were normalized blind to sample group (blind=TRUE) using the variance-stabilizing transformation (vst) in DESeq2.

### Gene ontology analyses of the transcriptome

For transcriptomic analyses, gene ontology (GO) overrepresentation analyses were performed using the clusterProfiler R package (v14.18.4; https://github.com/YuLab-SMU/clusterProfiler) with the *Mus musculus* annotation database org.Mm.eg.db. Lists of dysregulated transcripts were compared to a background comprising all detected transcripts after count filtering, i.e. mean count > 0 across samples excluding mitochondrially encoded and microglial genes associated with inhibitory neurons (as described for DESeq2 analysis). For GO overrepresentation analyses with P3 fractions, the custom background was defined with transcripts detected in P3 samples. For analyses with P3 and notP3 fractions the background was defined with P3 and notP3 samples. Ontology settings were set to “all” (CC, BP, MF) and p-values were calculated with Fisher’s exact test subjected to Benjamini-Hochberg (BH) multiple-testing correction. Significant terms were defined as FDR < 0.05. Complete GO overrepresentation result files are available as supplementary tables (**Table S6, S8, S14, S15, S20, S21**).

### Reference lists matching of transcriptomics data

To assess the overlap of the differentially regulated transcripts with previously published studies and databases such as SFARI (all database entries, downloaded on December 11^th^ 2025; [52]), the NDD Gene Hub (formerly Geisinger DBD Genes; Loss-of-Function table, downloaded on August 15^th^ 2025; [53]), sex chromosomal genes (extracted from GENCODE (GRCm39) using the R package “rtracklayer” [54] and imprinted genes [55] a gene-level approach was used to integrate the different datasets, i.e. transcripts with the same gene name were matched at the level of gene symbols. Published data from Donnard and colleagues were filtered for FDR < 0.01 [8] and from Suresh and colleagues for FDR < 0.05 [39] to define a significantly dysregulated subset for each dataset.

### MS sample preparation

#### Bulk samples

Lysates of tissue homogenate or unsorted (crude) synaptosomes (n_KO_ = 9 *Fmr1*^-/y^ and 6 *Fmr1^-/+^*, n_WT_ = 7 *Fmr1*^+/y^ and 8 *Fmr1*^+/+^) were prepared via an adapted suspension trapping protocol for bottom-up proteomic analyses, as previously reported [56]. In brief, all samples containing 25 µg total protein, were mixed with 2x lysis buffer (10% sodium dodecyl sulfate [SDS; Sigma-Aldrich; cat# L4509], 100 mM TRIS, pH 7.55 adjusted with phosphoric acid [H_3_PO_4_; Supelco, Bellefonte, PA, USA; cat# 49685]) in a 1:1 ratio and reduced using dithiothreitol in a final concentration of 20 mM for 10 min at RT. For cysteine alkylation, the samples were incubated with iodoacetamide in a final concentration of 50 mM for 30 min at RT in the dark. Then, lysates were acidified using H_3_PO_4_ in a final concentration of ∼1.2%. Binding/ wash buffer (90% methanol, 50 mM TRIS, pH 7.1 with H_3_PO_4_) was added in a 1:7 lysate-to-buffer ratio. The protein suspension was loaded onto the filter of an S-trap (size = micro; ProtiFi, Fairport, NY, USA; cat# C002-MICRO) by centrifugation for 20 s at 4,000 x g in 150 µl-steps. Trapped proteins were washed with 150 µl binding/ wash buffer four times. MS-grade trypsin (1 µg; Promega, Madison, WI, USA) was added in 60 µl of a digestion buffer (ammonium bicarbonate [ABC]; 50 mM; Sigma-Aldrich, St. Louis, MO, USA; cat# 09830) and digestion was carried out overnight at RT in a humidified chamber. For peptide elution, the filter was washed in three consecutive steps by centrifugation at 4,000 x g for 40 s with 40 µl digestion buffer and two 40 µl washes 0.2% formic acid (FA; Merck Millipore, Burlington, MA, USA; cat# 5.33002) in MS-grade water (H_2_O; VWR, Radnor, PA, USA; cat# 83645.29). Peptides were dried *in vacuo* and stored at -20°C until LC-MS analyses.

#### Sorted synaptosomes

We prepared 19 WT and 25 *Fmr1*-KO synaptosomes from NEX-Cre::SypTOM^+^ animals (n_WT_ = 8 *Fmr1*^+/y^ and 11 *Fmr1*^+/+^, n_KO_ = 12 *Fmr1*^-/y^ and 13 *Fmr1^-/+^*) as well as 24 WT and 20 *Fmr1*-KO synaptosomes from Gad2-Cre::SypTOM^+^ animals (n_WT_ = 10 *Fmr1*^+/y^ and 14 *Fmr1*^+/+^, n_KO_ = 6 *Fmr1*^-/y^ and 13 *Fmr1^-/+^*), similar to previous reports [3]. Sorted synaptosomes were filtered onto Whatman glass microfiber filters (Cytiva, Marlborough, MA, USA; cat# 1825-090) as described above, and stored at –80°C in LoBind Eppendorf tubes until further processing for bottom-up proteomics. The samples (2 million sorted particles) were thawed on ice and protein digestion was carried out using a digestion buffer containing trypsin (0.1 μg; Promega, Madison, WI, USA; cat# V5117) and rLysC (0.1 μg; Promega, Madison, WI, USA; cat# V1671) in 25 μl 50mM TEAB for each sample. Digestion was performed overnight (16-18 hours) at 37 °C, shaking at 300 rpm. 25 μl pure MS-grade acetonitrile (ACN; VWR, Radnor, PA, USA; cat# 83640.29) was then added to each tube and the samples were centrifuged at 16,000 × g for 10 min. A pipette tip containing C_18_ material (µZipTip, Merck Millipore, Burlington, MA, USA; cat# ZTC18M096) was washed once using 50% ACN in H_2_O. Using the ZipTips, the supernatants of the digest were filtered by centrifugation for 2 min at 2,000×g into MS vials (QuanRecovery, Waters, Milford, MA, USA; cat# 176004434). Then, 50 μl 50% ACN in H_2_O was added to each sample to eluate remaining peptides and the filtering process was repeated as just described. Peptides eluted into the MS vials were concentrated *in vacuo* at 45 °C until full dryness and stored at -20°C until LC-MS analysis.

### LCMS analysis

Dried peptides of sorted synaptosome preparations were resuspended in 8.5 µl reconstitution buffer (5% ACN, 95% H_2_O, 0.1% FA supplemented with indexed retention time peptides (iRT; Biognosys, Schlieren, Switzerland; cat# 1816351) in a 1:100 dilution. Dried peptides of homogenate and crude synaptosome preparations were reconstituted in 25 µl reconstitution buffer, respectively. Peptides were analyzed via a nano-UHPLC (U3000, Dionex, Sunnyvale, CA, USA) coupled to an Orbitrap Fusion Lumos tribrid mass spectrometer (ThermoFisher Scientific, Waltham, MA, USA) via a NanoFlex ion source, injecting 6 µl of the sorted synaptosomes and 1 µl of the bulk samples. Peptides were loaded onto a PepMap 100 C_18_ trap column (75 μm id × 2 cm length, 3 μm particle size; Thermo Fisher; cat# 164946) at 6 μl/min for 6 min with 2% ACN and 0.05% trifluoroacetic acid in H_2_O followed by separation on an C_18_ analytical column (75 μm id × 50 cm length, 1.7 μm particle size; CoAnn Technologies LLC, Richland, WA, USA; cat# HEB07505001718IWF) maintained at 55°C (nano column oven; Sonation). Analytes were separated in a 120 min non-linear gradient according to the published, paired LC-MS method [57], using mobile phase A (0.1% FA in H_2_O) and B (80% ACN, 20% H_2_O, 0.1% FA). Peptides were analyzed in positive polarity and data independent acquisition (DIA) mode. The DIA method defined MS1 scans followed by 40 DIA scans with optimized segment widths [57]. In brief, the MS method had the following settings. Full scan: orbitrap resolution = 120k, AGC target = 125%, mass range = 350-1650 m/z and maximum injection time = 100 ms. DIA scan: activation type = HCD, HCD collision energy = 27%, orbitrap resolution = 30k, AGC target = 2000% and maximum injection time = dynamic.

### MS data processing

#### Bulk samples

DIA raw files were processed with dia-nn (academic version 2.6.0) using a library-free approach [58]. The library was predicted using the *in silico* FASTA digest (Trypsin/P) option with the UniProtKB database for *Mus musculus* (UP000000598; downloaded June 11^th^ 2025), including the sequence for the Synaptophysin-tdTomato construct, the iRT peptides (Biognosys, Schlieren, Switzerland; cat# 1816351), and a database for common MS-contaminants as defined by dia-nn. Deep learning-based spectra and retention time prediction were enabled. Peptide length was set to 7-35 amino acids, missed cleavages to 2, and charge range to 1-6. N-terminal methionine excision was enabled. Methionine oxidation and N-terminal acetylation was set as variable modifications and Cysteine carbamidomethylation as fixed modification. The maximum number of variable modifications per peptide was one. Mass accuracy of 10 ppm, MS1 accuracy of 4 ppm, and scan window of 7 were selected. The precursor false-discovery-rate (FDR) level was set to 0.01. Protein inference was performed using (species-specific) genes including the protein inference option. The match-between-runs option was disabled.

The dia-nn report table and the respective FASTA files were imported into the statistical computing software R (v.4.3.3) and analyzed using the MSDAP R package (v 1.2.1; [59]). Contaminants, iRT peptides and keratins were discarded. MSDAP performed protein roll-up and computed label-free intensities at the protein level (minimum of one peptide per protein, maxLFQ algorithm). Normalization was performed using robust linear regression normalization (“rlr”) as sourced from the msqrob R package implemented in MSDAP. To compare WT and *Fmr1*-KO animals in CX homogenate and synaptosomes, differential protein abundance analyses were conducted using a peptide-level linear mixed effect model (“msqrob”), adding “litter” as a random variable from the sample metadata. Analyses were performed separately for hemizygous (*Fmr1*^-/y^ vs *Fmr1*^+/y^; males) and heterozygous (*Fmr1*^-/+^ vs *Fmr1*^+/+^; females) mice. Multiple testing correction was performed according to the Benjamini–Hochberg method with an FDR cut-off below 0.01. Significant hits quantified with less than 2 peptides per protein in the individual comparisons were discarded.

#### Sorted synaptosomes

DIA raw files were processed with dia-nn (academic version 2.3.0) using a library-free approach [58]. The library was predicted using the *in silico* FASTA digest (Trypsin/P) option with the UniProtKB database for *Mus musculus* (UP000000598; downloaded June 11^th^ 2025), including the sequence for the Synaptophysin-tdTomato construct, the iRT peptides, and a database for common MS-contaminants as defined by dia-nn. Peptide length was set to 7-35 amino acids, missed cleavages to 2, and charge range to 1-6. N-terminal methionine excision was enabled. Methionine oxidation was set as variable modification and Cysteine carbamidomethylation as fixed modification. The maximum number of variable modifications per peptide was one. Mass accuracy of 10 ppm, MS1 accuracy of 4 ppm, and scan window of 7 were selected. The precursor FDR level was set to 0.01. Protein inference was performed using (species-specific) genes including the protein inference option. The match-between-runs option was enabled. The dia-nn report table and the respective FASTA files were imported into the statistical computing software R (v.4.3.3) and analyzed using the MSDAP R package (v 1.2.1; [59]). MSDAP performed protein inference and label-free quantification at the protein level (minimum of one peptide per protein, maxLFQ algorithm). Normalization was performed using robust linear regression normalization (“rlr”) as sourced from the msqrob R package implemented in MSDAP. To compare WT and *Fmr1*-KO animals, differential protein abundance analyses were conducted using the “msempire” algorithm as implemented in MSDAP. Analyses were performed separately for hemizygous (*Fmr1*^-/y^ vs *Fmr1*^+/y^; males) and heterozygous (*Fmr1*^-/+^ vs *Fmr1*^+/+^; females) mice. Multiple testing correction was performed according to the Benjamini–Hochberg method with an FDR cut-off below 0.05. Significant hits quantified with less than 2 peptides per protein in the individual comparisons were discarded.

To assess purity of the synaptic preparation, the sorted fraction (P3) was compared over the control fraction (notP3) of WT excitatory and WT inhibitory synaptosomes. To assess cell type specificity, sorted fractions (P3) of inhibitory and excitatory WT synaptosomes were contrasted. Differential protein abundance analyses were conducted using a peptide-level linear mixed effect model (“msqrob”), adding “litter”, “sex”, and “animal” as a random variable from the sample metadata. Multiple testing correction was performed according to the Benjamini-Hochberg method with an FDR cut-off below 0.05.

Result files of all individual DAAs are available as supplemental tables. All data preparation was performed using the tidyverse- and visualization using the ggplot2-or pheatmap-packages in R.

### Reference lists matching and enrichment analyses of proteomics data

To assess the overlap of the differentially regulated proteins with previously published studies and databases such as SFARI (all database entries, downloaded on December 11^th^ 2025; [52]) or the NDD Gene Hub (formerly Geisinger DBD Genes; Loss-of-Function table, downloaded on August 15^th^ 2025;[53]), a gene-level approach was used to integrate the different datasets, i.e. proteins with the same gene name were aggregated and matching was performed at the level of gene symbols. Published data from Louros and colleagues were filtered for p-values below 0.05 to define a dysregulated subset [33]. P17-data from Tang and colleagues were filtered for BH-corrected p-values below 0.05 to define a dysregulated subset [34]. Results from Gao and colleagues were filtered for BH-corrected p-values below 0.05 and fold-changes as defined by the authors [60]. Enrichment of the selected reference list was tested against the experimentally detected protein background using one-sided Fisher’s exact tests (one-sided, alternative = "greater"), with Benjamini–Hochberg correction for multiple testing (BH−adjusted p < 0.05, ** p < 0.01, *** p < 0.001). For directionality analyses, only significant proteins with valid fold changes in either Gad2^+^, Nex^+^ or both synapse types were considered and classified as concordantly regulated (same fold change sign) or reciprocally regulated (opposite signs). Fold changes of significant entries were retained without averaging across duplicate background entries; genes with conflicting direction assignments among multiple significant entries were excluded from direction-based tests. Overrepresentation of reference list entries was then tested within concordant, reciprocal, and individual direction subclasses as described above.

### Gene ontology analyses of proteomics data

For all proteomics analyses, gene ontology overrepresentation analyses were performed using the clusterProfiler R package (v4.10.1; https://github.com/YuLab-SMU/clusterProfiler). The annotation file of genes from *Mus musculus* was accessed from OrgDB using the identifier AH114085 (NCBI gene ID based annotations, org.Mm.eg.db.sqlite). Gene lists of dysregulated proteins or functional protein clusters were compared to the individual, experimentally detected protein background (i.e. proteomics of CX homogenate, CX unsorted synaptosomes, or synaptic (P3) fractions) using Fisher’s exact tests and the ontology setting “all” (GOCC, GOMF, and GOBP). After applying Benjamini-Hochberg correction, an FDR cut-off below 0.05 was defined. Enrichment results were then merged and plotted by gene count for visualization of the top terms ranked by FDR according to reported condition (see Figure). Complete result files, beyond the plotted top-terms, are available as supplemental tables (**Tables S2, S4, S10, S12, 18, 19, 22**).

### Integration of transcriptomic and proteomic datasets

Data integration was performed at gene-level. To circumvent multiple entries due to Protein IDs with identical gene names, merging rules were the following: match mRNA to matching protein with (1) significant regulation, (2) if not significant - with most peptides used in comparison, (3) if all significant - use the one with most peptides. Overall comparative analyses of mRNA and protein data were performed using average log_2_ scaled DESeq2 normalized counts or protein intensities. Pearson correlation coefficients were calculated using pair-wise complete information of mRNA-protein matches. *Fmr1*-KO induced dysregulation was investigated at log_2_ fold change level for both molecules.

### Western Blot

Cortex homogenates were prepared by Dounce homogenization in PBS with Protease inhibitor cocktail III (1:750, Calbiochem; EMD, Merck Millipore, Burlington, MA, USA; cat# 539134-10ml) from one Cortex half of 7-9 week-old animals with different *Fmr1* genotypes. After protein determination we added NuPAGE 4x LDS Sample buffer (ThermoFisher, Waltham, MA, USA; NP0007), 10x reducing agent (ThermoFisher, Waltham, MA, USA; cat# NP0009) and water to get 400 µl lysate in 1xLDS/1x reducing agent with equal total protein concentration for the different genotypes. The samples were heated to 95°C for 5 min and 5 µg or 20 µg were separated on NuPAGE 4-12% Bis-TRIS gels (ThermoFisher, Waltham, MA, USA; cat# NP0321) with 3-(N-morpholino)propanesulfonic acid (MOPS) running buffer (ThermoFisher, Waltham, MA, USA; cat# NP0001) and blotted. Total protein was visualized by staining with Ponceau S solution (ThermoFisher, Waltham, MA, USA; cat# A40000279). After blocking with PBS Intercept blocking buffer (Licor Biosciences, Lincoln, NE, USA; cat# 927-70001) primary antibody (FMRP 1:2,000, Merck Burlington, MA, USA; cat# ZRB1081) was applied overnight at 4°C in PBS-Intercept blocking buffer/PBS 0.2% Tween-20 1+1 (PBST-Odyssey), IRDye800-coupled secondary antibody was used 1:20,000 and applied for 1 h at room temperature prior to scanning on an Odyssey F imager with Licor acquisition 2.4 software (Licor Biosciences, Lincoln, NE, USA).

### Immunohistochemistry

Brain hemispheres stored in 30% sucrose as described above were subjected to cryotome sectioning for 40 μm brain sections. Sections were washed 3 times in PBS and permeabilized overnight with 0.5% Triton X-100 in blocking buffer (BB; 4% goat serum in PBS pH 7.4). Immunostaining was performed by incubating samples with primary antibodies (gp-anti-vGAT 1:1000 SySy, Göttingen, Germany; cat# 131004; rb-anti-vGLUT1 1:2000 SySy, Göttingen, Germany; cat# 135303; rb-anti-FMRP, Merck Burlington, MA, USA; cat# ZRB1081; gp-anti-NeuN 1:500 SySy, Göttingen, Germany; cat# 266004) in BB overnight at 4 °C, followed by 4 times 10 min washes with PBS and a 4 h incubation with DAPI and gt-anti-rb or -gp secondary antibodies coupled to Alexa-488 or Alexa-647 (ThermoFisher Scientific, Waltham, MA, USA) in BB at RT. Sections were washed with PBS and mounted with AquaPolymount. Fluorescence imaging was performed with a LSM780 laser scanning confocal microscope (Zen10 software, Zeiss, Jena, Germany) using a Plan-Apochromat 40x/NA 1.4 oil DIC M27 objective or 20x and appropriate excitation laser lines and spectral detection windows. To assess correct synapse labelling, spatial colocalization between Nex-SypTOM or Gad2-SypTOM and the synaptic markers vGLUT1 and vGAT was evaluated using the “Coloc2” plugin in Fiji/ImageJ, applying identical settings for all images (maximum intensity projection). Intensity correlation between the two channels (X-SypTOM vs synaptic marker) was calculated using Pearson’s correlation. For each genotype, one section with six fields of view was included. As additional control for random colocalization, the same analysis was performed with one channel rotated by 90°.

## Results

### Profiling Fmr1-KO synapses with cell type-specificity using FASS, RNA-Seq and DIA-MS

To investigate the effect of FMRP loss at synaptic resolution, we optimized our previously established FASS pipeline [3, 2] to enrich and analyze cell type-specific synaptosomes from the cortex of *Fmr1*-KO mice (*Fmr1*^-/y^ or *Fmr1*^-/+^) and wildtype litter mates (WT; *Fmr1*^+/y^ or *Fmr1*^+/+^) at postnatal week 7. We used Cre-inducible knock-in mice to express the synaptic vesicle protein synaptophysin fused to the fluorescent marker tdTomato (SypTOM) to label the presynaptic compartment (**Figure 1a**). To distinguish excitatory, glutamatergic and inhibitory, GABAergic neurons, Nex-Cre (Nex^+^; [41]) and Gad2-Cre (Gad2^+^; [40]) driver mouse lines were used. We developed breeding strategies to generate littermate animals carrying the *Fmr1*-KO allele or the WT allele with labeled excitatory or inhibitory synapses. One cortical hemisphere was processed for imaging, the other hemisphere was used to prepare and sort 2 million excitatory or inhibitory synaptosomes (P3 fraction) from individual WT or *Fmr1*-KO animals (7 weeks) for downstream RNA-Seq or DIA-MS. We collected input fractions (cortex homogenate and unsorted synaptosomes) and sorted control fractions deenriched for the fluorescent population (notP3 fraction) for quality control of the *Fmr1*-KO model (**Figure S1; Tables S1-S4**) as well as synaptic purity and cell type-specificity of our preparations (**Figure S2; Tables S5-S12**). Following FASS, we obtained high purity for both synapse types and conditions (**Figures 1b, S2, Tables S1-S12**). We observed a pronounced average enrichment to 94% and 91% in excitatory and inhibitory synapses, respectively (**Figure 1b**). Since alterations in synapse number and/or morphology have been described in *Fmr1*-KO mice [61, 22–24], we inspected our FASS results and noted no systematic differences between WT and KO synapses.

**Figure 1:**
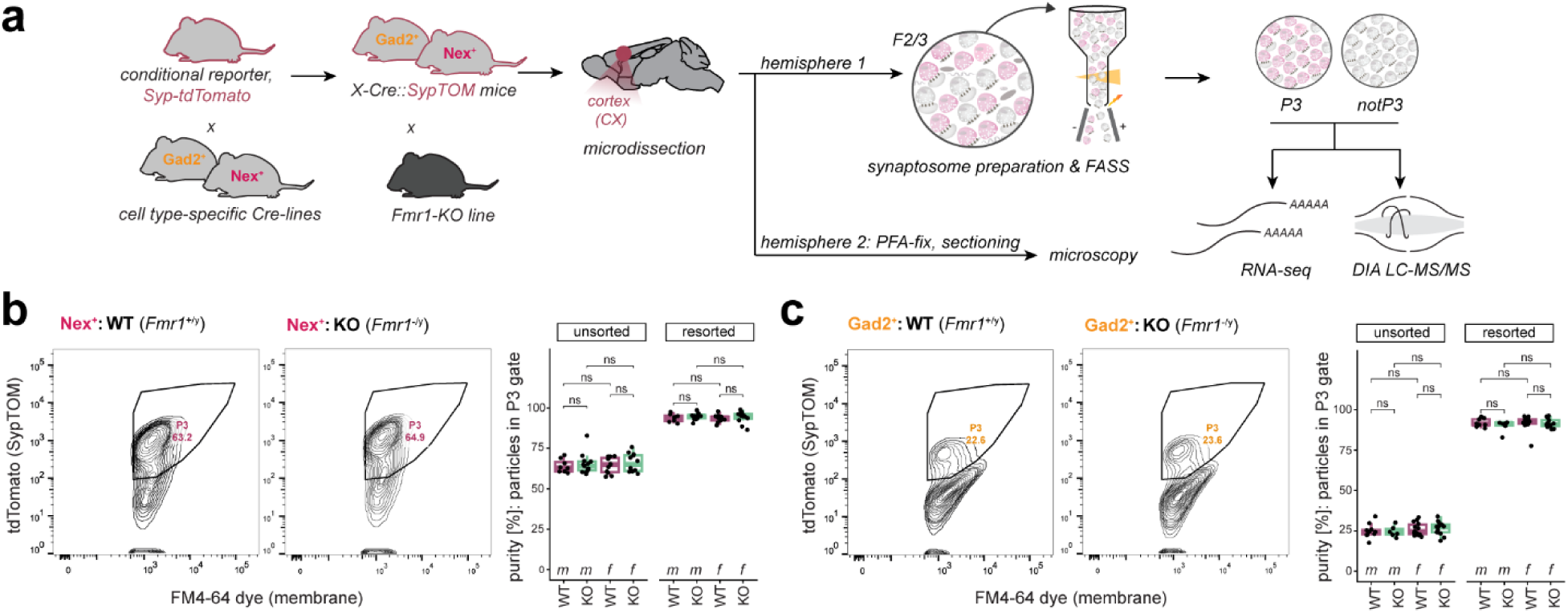
A pipeline for in-depth characterization of *Fmr1*-KO synapses with cell type-specificity. **a)** Fluorescence-activated synaptosome sorting (FASS) methodology and its required steps to assess the molecular diversity of cortical synapses from *Fmr1*-KO mice compared to WT litter mates (7 weeks of age). **b,c)** FASS contour plots show the relative density of the targeted SypTOM^+^ synaptic population (P3 gate) from cortical synaptosomes of WT (*Fmr1*^+/y^) or KO (*Fmr1*^-/y^) mice, for selected sorts of Nex^+^ (**b**) and Gad2^+^ (**c**) samples. Quantification of relative enrichments for both synapse types and sexes are summarized in boxplots by visualization of the synaptic population (P3 gate) of the first sorting run (unsorted) and its subsequent re-loading (resorted) to control for proper enrichments. During FASS, no significant differences between the populations of sorted particles have been observed for WT (*Fmr1*^+/y^ or *Fmr1*^+/+^) and *Fmr1*-KO (*Fmr1*^-/y^ or *Fmr1*^-/+^) animals (unpaired, two-sided Welch’s t-test). ns: not significant.

### The excitatory and inhibitory transcriptome of Fmr1-KO synapses

As FMRP is a prominent RNA-binding protein, we first queried whether the synaptic transcriptome is altered in *Fmr1*-KO (*Fmr1^-/y^*) synapses in males (**Figure 2a**). Using RNA-Seq, we identified ∼7,000 transcripts in both *Fmr1^+/y^* (WT) and *Fmr1^-/y^* animals and in both synapse types (**Figure 2b**). Principal component analysis (PCA) revealed distinct *Fmr1^-/y^* and *Fmr1^+/y^* clusters in excitatory and inhibitory synaptosomes, indicating a remodelling of the transcriptome (**Figure 2b**). To identify dysregulated mRNA candidates, we performed differential expression analyses (DEA), revealing changes in both synapse types (**Figure 2c, Table S13**). Excitatory synapses showed 109 and 50 significantly up-or downregulated transcripts, respectively. Inhibitory synapses exhibited a similar number of dysregulated mRNAs: 99 up- and 56 downregulated. Gene ontology (GO) analysis revealed significant overrepresentation of synaptic terms, including postsynaptic density, regulation of synaptic structure and asymmetric synapse for dysregulated excitatory synapse transcripts (**Figure 2d, Table S14**). Transcripts that gave rise to these overrepresented groups were associated with vesicle trafficking (*Iqsec1)*, synaptic plasticity (*Gsk3b*, *Dgkb)* or microtubule components (*Map2, Tubb5)*. In inhibitory synaptosomes, downregulated transcripts showed a significant overrepresentation for dopamine receptor binding including transcripts encoding the G-protein coupled proteins *Gnai1* and *Gnas,* and for phosphatidylinositol-4-phosphate binding including the GTPase-activating protein *Dab2ip*. We noted that 16% of dysregulated transcripts of each synapse type were ASD-risk genes (Simons Foundation Autism Research Initiative. SFARI; [52]) and ∼10% were annotated as NDD-risk genes (NDD Gene Hub, [53] (**Figure 2e, Table S13**). In addition, just ∼15% of dysregulated transcripts in inhibitory and ∼19% in excitatory synapses were established FMRP targets [15] indicating that the majority of dysregulated transcripts are not previously identified as FMRP-bound targets. To investigate if the synapse type-specific dysregulation is also reflected by somatic transcriptomic alterations upon *Fmr1*-KO, we integrated our synapse type-specific transcriptomic dataset with a single-cell RNA-Seq (scRNA-Seq) dataset of cortical excitatory and interneuron somata from *Fmr1*-KO mice [8]. In inhibitory synapses only ∼18% (29/155) and in excitatory synapses ∼24% of dysregulated transcripts (38/159) showed an expression pattern similar to that observed in somatic scRNA-Seq data of cortical excitatory and interneurons, respectively, with an overall low correlation (R_Nex+_ = 0.13, R_Gad2+_ = 0.00029; **Figure S3a**). This indicates that the majority of synapse type-specific transcriptomic dysregulation observed here is not a passive reflection of somatic alterations and that the loss of FMRP leads to distinct subcellular transcriptomic landscapes.

**Figure 2:**
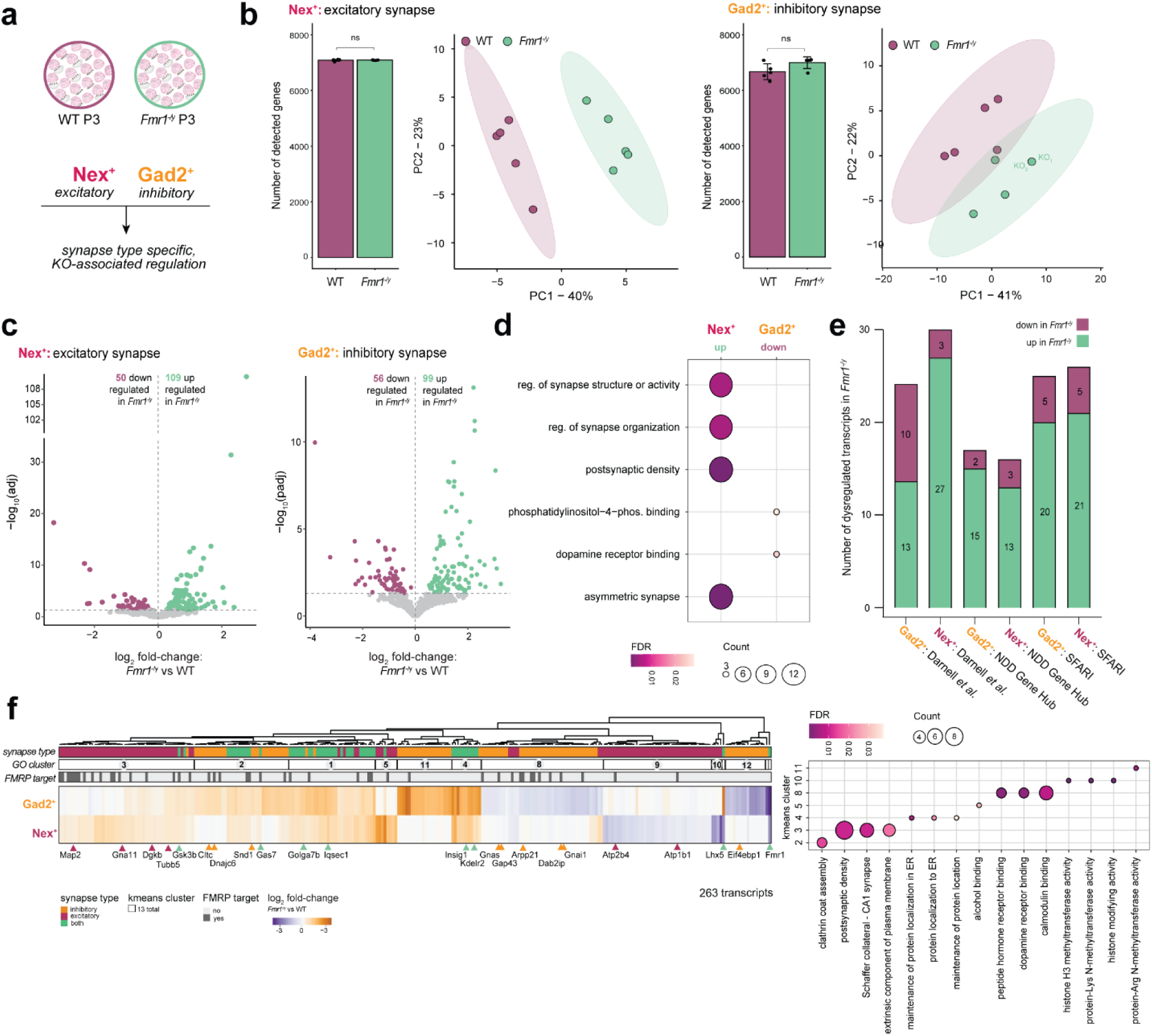
Cell type-specific mRNA signatures of *Fmr1*-KO (*Fmr1*^-/y^) synapses. **a)** Analyses strategy to assess *Fmr1*-KO induced mRNA changes within the cell type-specific synaptic compartment. **b)** Barplots show quantification of ∼7,000 detected genes in both Nex^+^ (left) and Gad2^+^ (right) sorted synaptosomes after filtering (2 million particles, see methods) with no significant difference for *Fmr1*^-/y^ and WT (unpaired, two-sided Welch’s t-test). PCA indicates clear separation of WT and *Fmr1*^-/y^ in Nex^+^ or Gad2^+^ sorted synaptosome transcriptomes (top 500 most variable genes; overlapping Gad2^+^ KO replicates are labelled). ns: not significant. n = 5 animals per condition. **c)** Volcano plots show results of differential expression analyses for excitatory (left) and inhibitory (right) synapses (FDR < 0.05). **d)** Dot plots show significant results of GO analyses of significantly up- and downregulated genes, respectively (GO terms from MF, BP, CC). **e)** Number of significantly dysregulated transcripts in *Fmr1*^-/y^ mice annotated as established FMRP mRNA targets [15], NDD-risk genes (NDD Gene Hub, [53]) and ASD-risk genes (SFARI database, [52]) for both synapse types. **f)** Heatmap of log_2_ fold changes for *Fmr1*^-/y^ vs WT of significantly dysregulated transcripts upon *Fmr1*-KO in both synapse types (left). Significant regulation according to synapse type, results of a segmentation analysis (GO clusters, n = 13), and FMRP mRNA target status [15] are conditionally highlighted below the dendrogram. The top 3 results of GO analyses for clusters yielding significant terms (clusters 2-5, 8, 10-11) are summarized in the dot plot (right) (GO terms from MF, BP, CC; FDR < 0.05).

To investigate if the dysregulation of transcripts was shared or specific for the respective synapse type, we performed unsupervised, hierarchical clustering (**Figure 2f**). Unsupervised clustering yielded 13 clusters and indicated that 2/3 of significantly dysregulated transcripts were synapse type-specifically up-(Nex^+^: clusters 3, 5, Gad2^+^: clusters 2, 11) or downregulated (Nex^+^: clusters 9, 10, Gad2^+^: clusters 8, 12, 13). The remaining 1/3 showed a common dysregulation pattern across synapse types (clusters 1, 2, 4, 6) where almost all transcripts were upregulated. Only one gene (cluster 7) showed significant reciprocal dysregulation between synapse types: the transcription factor *Lhx5* was upregulated in inhibitory and downregulated in excitatory synapses. To understand which functional pathways were affected upon *Fmr1*-KO in a synapse type-dependent and -independent manner, we performed GO analyses for the individual clusters yielding significant GO overrepresentations for 7/13 clusters (**Figure 2f**, **Table S15**). This analysis revealed that similar processes important for synaptic function were affected: commonly upregulated transcripts were essential for protein localization and lipid metabolism in ER (cluster 4: *Kdelr2*, *Insig1*), synaptic vesicle recycling/transmission (cluster 2: *Gas7*), and cell adhesion (cluster 1: *Golga7b*). Excitatory synapses exclusively upregulated transcripts were related to synaptic organization (cluster 3; as observed for **Figure 2d**) while exclusively downregulated transcripts included ion channel subunits essential for maintaining cation homeostasis (cluster 9: *Atp1b1*, *Atp2b4*). Inhibitory synapses additionally showed an exclusive upregulation of synaptic vesicle recycling components (cluster 2: *Cltc*, *Dnajc6*) and downregulation of transcripts required for diverse synaptic mechanisms ranging from calmodulin interactions and receptor binding (cluster 8: *Gap43*, *Arpp21*, *Gnai1*, *Gnas*) to local translation regulation (cluster 12: *Eif4ebp1*). These synapse type-specific transcriptomic signatures are not due to the synapse type origin but rather induced by FMRP loss (**Figure S2e, Tables S7-S13**).

### The excitatory and inhibitory proteome of Fmr1-KO synapses

Our proteomic profiling of the WT and *Fmr1*-KO (*Fmr1^-/y^*) synapses reliably quantified over ∼4,000 proteins for all FASS preparations of both excitatory and inhibitory synaptosomes (**Figure 3a**). Unlike the transcriptome, PCA of the proteome revealed less pronounced KO-associated separation between WT and *Fmr1*-KO excitatory and inhibitory synapses (**Figure 3a**). MS quantification of FMRP itself resulted in detection in both glutamatergic and GABAergic WT synapses at low levels and its absence in KO samples (**Figure 3a**). Differential abundance analyses (DAAs) of the synaptic proteome of WT and *Fmr1*-KO (*Fmr1^-/y^*) mice yielded 87 and 67 significantly dysregulated proteins for excitatory and inhibitory synapses, respectively (**Figure 3b**, **Table S16**). GO analyses for the individual synapse types (**Table S17**) revealed terms associated with synaptic functions, e.g. postsynaptic density for glutamatergic or structural constituent of the synapse for GABAergic synaptosomes, which all shared significant downregulation independent of synapse type (**Figure 3b**). These groups are reflected by protein candidates such as CTNNB1, SNAP25, GRIN1 for excitatory or SPTBN2 and SHANK1 for inhibitory synapses, respectively. For the top upregulated functions and pathways, we observed little overlap between synapse types. While proteins associated with kinase binding (e.g. MYH9, VDAC1, SRCIN1, MAP2, TUBA4A) or myelin (e.g. SYN1, GNAO1, PGAM1, VDAC1, COX5A) were enriched in excitatory synapses, proteins associated with mitochondrial terms (e.g. MRPL10, LRPPRC) and energy metabolism (e.g. GLS, OGDHL, SLC25A12) showed upregulation in inhibitory synapses.

**Figure 3:**
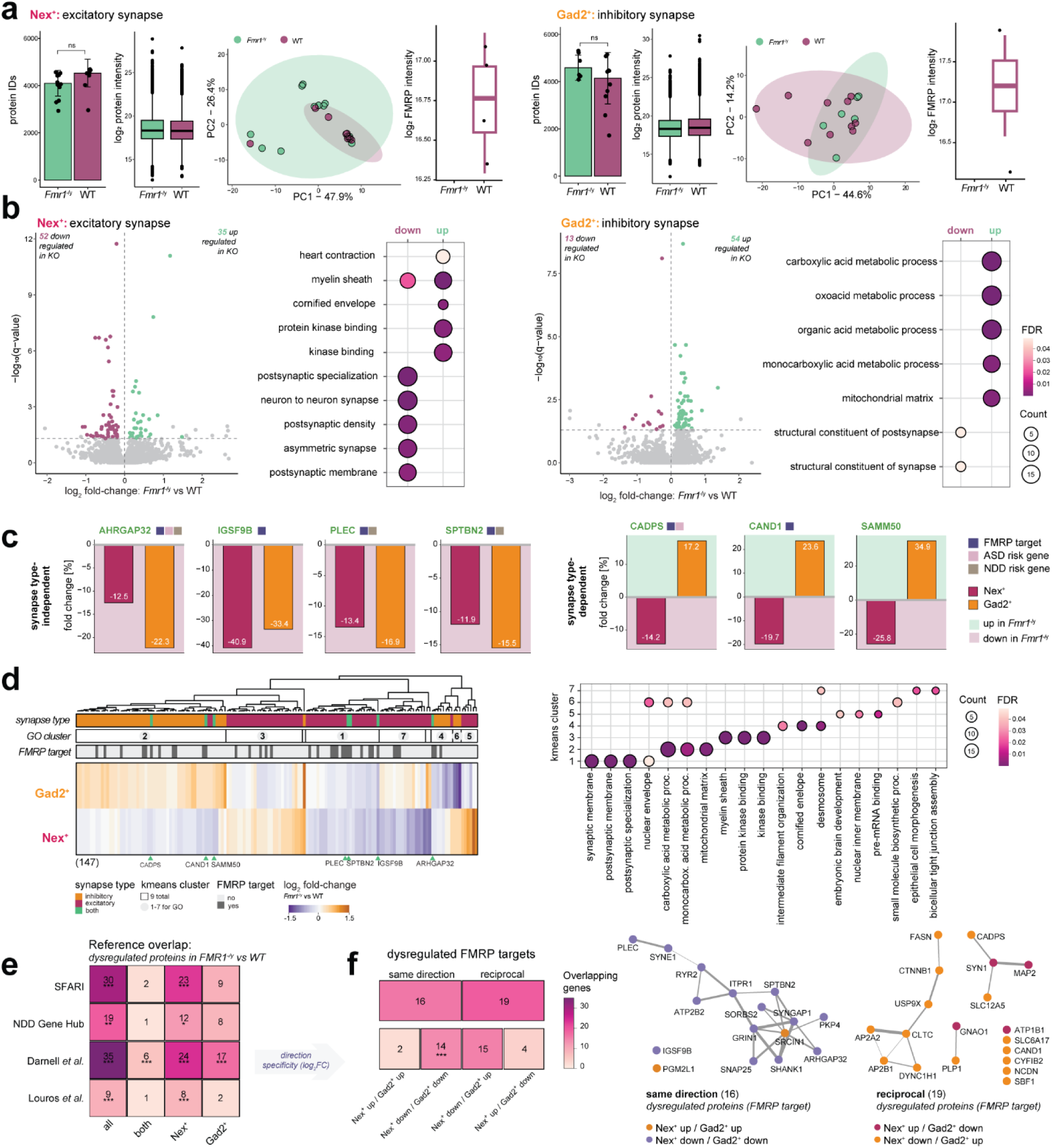
Cell type-specific protein signatures of *Fmr1*-KO (*Fmr1*^-/y^) synapses. **a)** Barplots show quantification of ∼4,000 proteins in the preparations of 2 million sorted synaptosomes of both Nex^+^ (left) or Gad2^+^ (right), with no significant difference for *Fmr1*^-/y^ and WT (unpaired, two-sided Welch’s t-test). High similarity is also observed for boxplots of overall protein intensities in both synapse types and conditions. PCA indicates no clear separation of WT and *Fmr1*-KO in Nex^+^ or Gad2^+^ sorted synaptosome proteomes (top 500 most variable proteins, log_2_ protein intensity). Boxplots display FMRP abundance at each synaptic compartment of WT mice and its absence at synapses of KO mice (log_2_ protein intensity). ns: not significant. **b)** Volcano plots show results of differential protein abundance analyses for excitatory (top) and inhibitory (bottom) synapses (**Table S16**). Both synapse types present significant abundance changes of the synaptic proteome (FDR < 0.05; Nex^+^: 87, Gad2^+^: 67 proteins). Dot plots summarize results of GO analyses (left; top-5, GOMF/BP/CC, FDR < 0.05), indicating few shared dysregulated (downregulated) pathways and different effects for upregulated ones. Dot size reflects the number of matched proteins and color indicates significance (FDR < 0.05). **c)** Barplots of candidates with significant abundance changes in both synapse types, highlighting synapse-type independent (top row) and dependent (bottom row) regulation (*Fmr1*^-/y^ vs WT fold change in percent). **d)** Heatmap of log_2_ fold changes for *Fmr1*^-/y^ vs WT of significantly dysregulated proteins upon *Fmr1*-KO in either (relaxed group) or both synapse types (left). Significant regulation according to subtype (synapse type), results of the segmentation analysis (GO cluster, n = 9), and FMRP mRNA target status [15] are conditionally highlighted below the dendrogram. The results of GO analyses for clusters yielding significant terms (clusters 1-7) are summarized in the dot plot (right), as described in b) (FDR < 0.05). **e)** Heatmap visualizes the overlap of the dysregulated subsets introduced in d) with known ASD-risk genes (SFARI database, [52]), risk genes of neurodevelopmental disorders (NDD Gene Hub, [53]), previously reported FMRP mRNA targets [15], and a previously published proteomics analyses of *Fmr1*^-/y^ mice using unsorted hippocampal synaptosomes [33]. The number of overlapping candidates is labelled in the cell and significance is highlighted using asterisks (one-sided Fisher’s exact test; BH-adjusted * p < 0.05, *** p < 0.001). **f)** Heatmap shows direction-pattern association of FMRP targets within significant proteins based on their log_2_ fold change. Analysis performed and labelled as described in e). Candidates changing in the same direction/ cell-type independent (left) or reciprocally/ cell type-dependent (right) are visualized as protein interaction networks (StringDB, *Mus musculus*). Node color indicates direction of dysregulation, edge width reflects the interaction score (StringDB combined score: 0.4-1).

We found seven candidates that were significantly dysregulated in both Nex^+^ and Gad2^+^ synaptosomes. When focusing on their dysregulation patterns, we found two different modes: a set of four proteins showed a shared downregulation (ARHGAP32, IGSF9B, PLEC, SPTBN2), while three candidates were similarly upregulated in inhibitory and downregulated in excitatory synapses (CADPS, CAND1, SAMM50) (**Figure 3c**). A majority (6/7) of these hits are previously reported FMRP target mRNAs [15], 2/7 encoded by ASD-risk genes [52], and 3/7 have links to NDD-associated missense mutations [53]. The only previously unreported candidate, SAMM50, a core component of the mitochondrial SAM complex [62], fits with reports of altered mitochondrial homeostasis in FXS [63–65].

To identify candidates with less stringency (relaxed group), we performed unsupervised, hierarchical clustering of proteins significantly changed in either excitatory or inhibitory, or both subtypes (**Figure 3d**). We found that most protein abundance changes occurred in an opposing manner in excitatory and inhibitory synapses. Subsequent GO analyses of the generated clusters yielded significantly enriched terms for 7/9 clusters (**Figure 3d**; **Table S18**). The segmentation analysis underscored the common downregulation of synaptic candidates (cluster 1), represented by GO terms such as synaptic membrane and postsynaptic specialization. For the larger group of opposingly regulated candidates, we confirmed previous mitochondrial or TCA-cycle-associated (cluster 2, Gad2^+^-KO up) and kinase-binding hits (cluster 3, Nex^+^-KO up). We further identified other strongly reciprocally dysregulated protein functions, such as organization of intermediate filaments (cluster 4, Nex^+^-KO up), mRNA binding and brain development (cluster 5, Nex^+^-KO up), small molecule biosynthesis (cluster 6, Gad2^+^-KO up), and tight junction assembly (cluster 7, Gad2^+^-KO slightly up). In mixed preparations such as cortex homogenates, there was less or a “diluted” dysregulation for the synapse type-specific candidates - reciprocal ones in particular - hence masking the adaptation and hindering detection (**Figure S4, Table S17**). To rule out the contribution of cell type-enriched protein abundances in the “baseline” synaptic proteomes, we integrated the results of a DAA comparing excitatory and inhibitory WT synaptosomes (**Figure S2e; Table S18, S19**) to our disease-associated findings and found no systematic pattern for cell type-preferential dysregulation, linking the reciprocal changes to the disease-phenotype rather than the cell type of origin. To assess whether the observed local dysregulation in Nex^+^ and Gad2^+^ synapses is a correlated cell-wide phenomenon, we compared to RiboTag-Seq data of excitatory and inhibitory neurons [39] and found just a modest correlation (R_Nex/Camk2a_ ∼ 0.14; R_Gad2/PV_ ∼ 0.39; **Figure S3b**).

There was a general overrepresentation of candidates from a previous proteomic analysis of bulk synaptosomes [33], ASD/NDD-risk genes, as well as previously established FMRP mRNA targets for the dysregulated proteins in both synapse types (35/147; **Figures 3e, S5a**). The overlapping FMRP targets represented proteins that exhibited reciprocal (19/35 proteins) and uni-directional (16/35 proteins) abundance changes (**Figure 3f**). Of note, proteins that exhibited a shared downregulation in Gad2^+^- and Nex^+^-synaptosomes constituted the only subset that showed significant overlap with FMRP targets (**Figure 3f**), comprising predominantly synaptic proteins (SHANK1, SNAP25, SYNGAP1) and proteins linked to intracellular Ca^2+^ signalling or transport (RYR2, IPTR2, ATP2B2).

### Loss of FMRP affects different mRNAs and proteins across synapse types

Are the regulated mRNA and protein populations correlated? Overall, we observed a moderate correlation between the detected level of a given mRNA and its protein for both *Fmr1^-/y^* or WT excitatory and inhibitory synapses (**Figure 4a**, R_Nex_ ∼ 0.38, R_Gad2_ ∼ 0.35) consistent with previous reports of synaptic protein-mRNA relationships [1, 4]. Strikingly, for candidates identified in the DAAs of both transcriptomics and proteomics (**Figure 2,3**), there was not much concordant regulation of mRNA and proteins (**Figure 4b,c**). We thus focused our analysis on molecules that showed significant dysregulation in the individual DAA by filtering mRNA-protein pairs regulated for either mRNA or protein (n_Nex_ = 144, n_Gad2_ = 127, **Figure 4b**). We compared the log_2_ fold changes of these transcripts and proteins and found mRNA-protein discordance. Separately analyzing pre-defined FMRP targets also did not reveal coordinated mRNA-protein regulation (**Figures 4b, 2e, 3e,f**). Generally, although only six candidates showed shared significant dysregulation (n_Nex_ = 5; n_Gad2_ = 1), all have been previously associated with ASD or NDDs. Three of the commonly dysregulated mRNA-protein pairs displayed coordinated regulation of mRNA and protein levels, namely *Map2*/MAP2 and *Aldh7a1*/ALDH7A1 in excitatory as well as *Cltc*/CLTC in inhibitory synaptosomes (**Figure 4c**). Interestingly, *Cltc*/CLTC and *Map2/*MAP2 showed a similar dysregulation pattern in RiboTaq-Seq data in Camk2a^+^ neurons ([39]; **Figure S3b,c**). The other three dysregulated transcript-protein ratios (*Sod1*/SOD1, *Etfb*/ETFB, *Atp1b1*/ATP1B1) observed in excitatory synaptosomes were discordant (**Figure 4c**).

**Figure 4:**
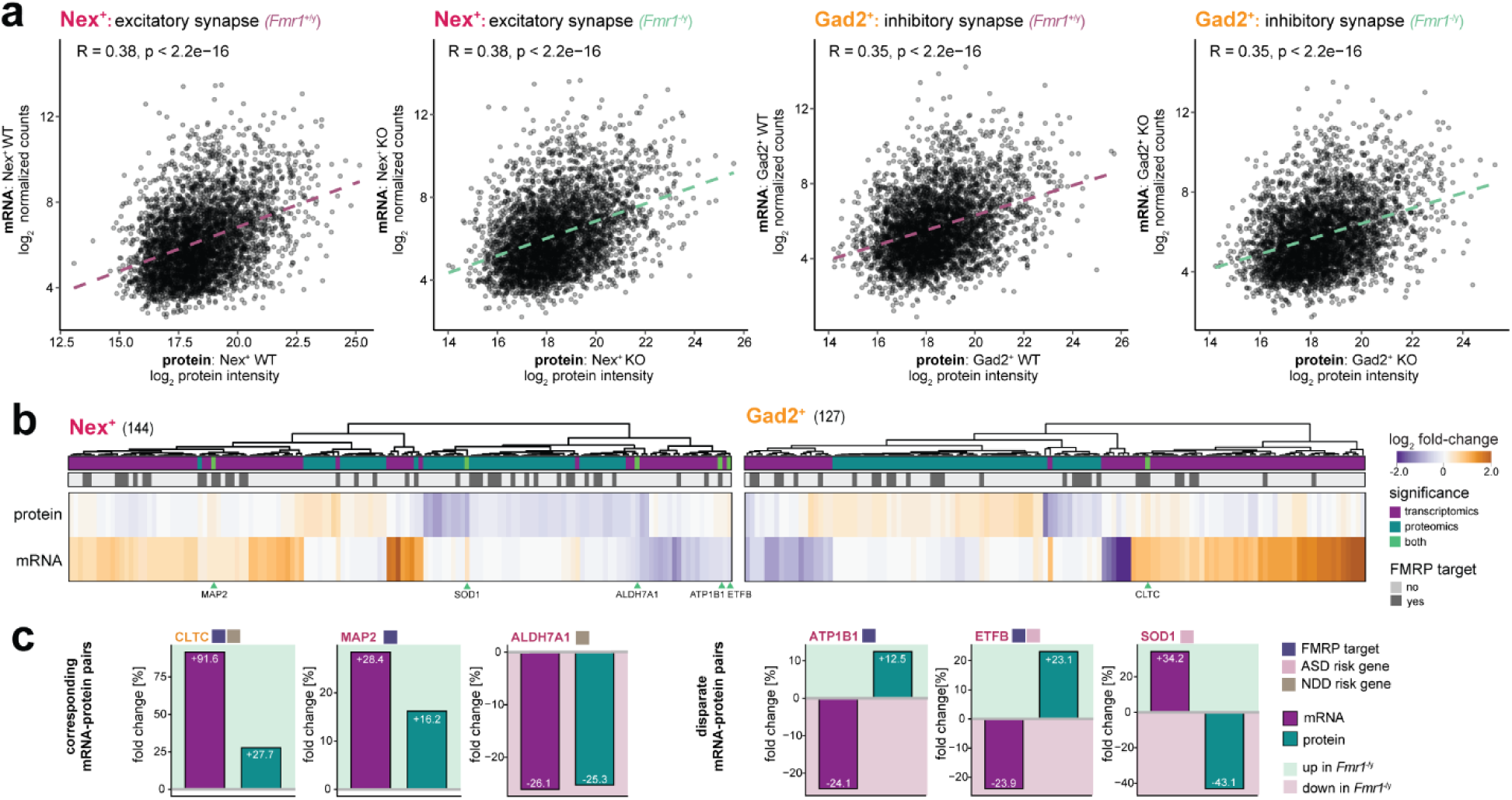
Differences in *Fmr1*-KO (*Fmr1*^-/y^) associated mRNA and protein dysregulation at excitatory and inhibitory synapses. **a)** Scatterplots show correlation of mean abundances of mRNAs (log_2_ DESeq normalized counts) and their respective protein (log_2_ protein intensity) for WT (*Fmr1*^+/y^) and *Fmr1*-KO (*Fmr1*^-/y^) animals of excitatory (left) and inhibitory (right) synapses, quantified by Pearson’s correlation and summarized with a linear regression fit. **b)** Heatmaps show log_2_ fold changes (*Fmr1*-KO vs WT) of significantly dysregulated mRNAs and proteins. Significance of dysregulation at transcript and/or protein level and reported FMRP mRNA target status [15] are conditionally highlighted below the dendrogram. Candidates significantly changed in both omics-regimes are labelled. **c)** Barplots of the shared dysregulated mRNA-protein pairs show corresponding and disparate regulation using the fold change (in percent). Color of the candidate name indicates dysregulation observed in Nex^+^ or Gad2^+^ synaptosomes. Functional annotation with respect to previously reported FMRP targets or ASD/ NDD-risk genes (NDD gene hub, SFARI database) is provided next to the gene name.

### Dysregulation of protein degradation machinery in Fmr1-KO excitatory and inhibitory synapses

Previous studies of *Fmr1-*KO synapses have identified alterations in both protein synthesis and degradation machineries [34, 27, 33]. We observed an increased abundance of ribosomal proteins (RPs) in excitatory *Fmr1*-KO synapses (also observed by [27]) and a decreased abundance of RPs in inhibitory synapses (**Figure 5a**). Focusing on components of the ubiquitin-proteasome system (UPS), we noted an enrichment of proteasome subunits (PSMA-D) in both synapse types, (see also [33]) (**Figure 5b**). As we also detected synapse type-specific alterations in individual UPS components (e.g. USP9X, CAND1), we next asked whether differential UPS target selection could contribute to the distinct pattern of proteomic remodeling observed in Nex^+^ and Gad2^+^ synaptosomes. The predicted ASD-risk gene CAND1 [66] caught our interest, as it is a key regulatory factor of Cullin-RING E3 ligase complexes (CRLs) [67, 68] that exhibited significant, but opposite patterns of dysregulation in both excitatory and inhibitory *Fmr1*-KO synapses (**Figure 5c**). Cullin proteins (CULs), reported to interact with CAND1 (**Figure 5d**), are localized at synapses (**Tables S9, S16**), but did not change upon FMRP loss. Integrating published CAND1/CUL interactomes [69] with our dysregulated proteins revealed overlapping candidates in a cell type-dependent manner (**Figure 5e**). While there was redirection of these interaction partners upon FMRP loss not always mirroring CAND1’s regulation pattern, they never showed the same direction of dysregulation in excitatory and inhibitory synapses. Of note, the identified CUL3-interactor CTNNB1 matched CAND1’s dysregulation. Further, it is part of a protein population that accumulates in proteasome inhibitor-treated, unsorted synaptosomes of hippocampi from *Fmr1*-KO mice, suggestive of a potential target of increased degradation in *Fmr1*^-/y^ synapses ([33]; **Figure S5**).

**Figure 5:**
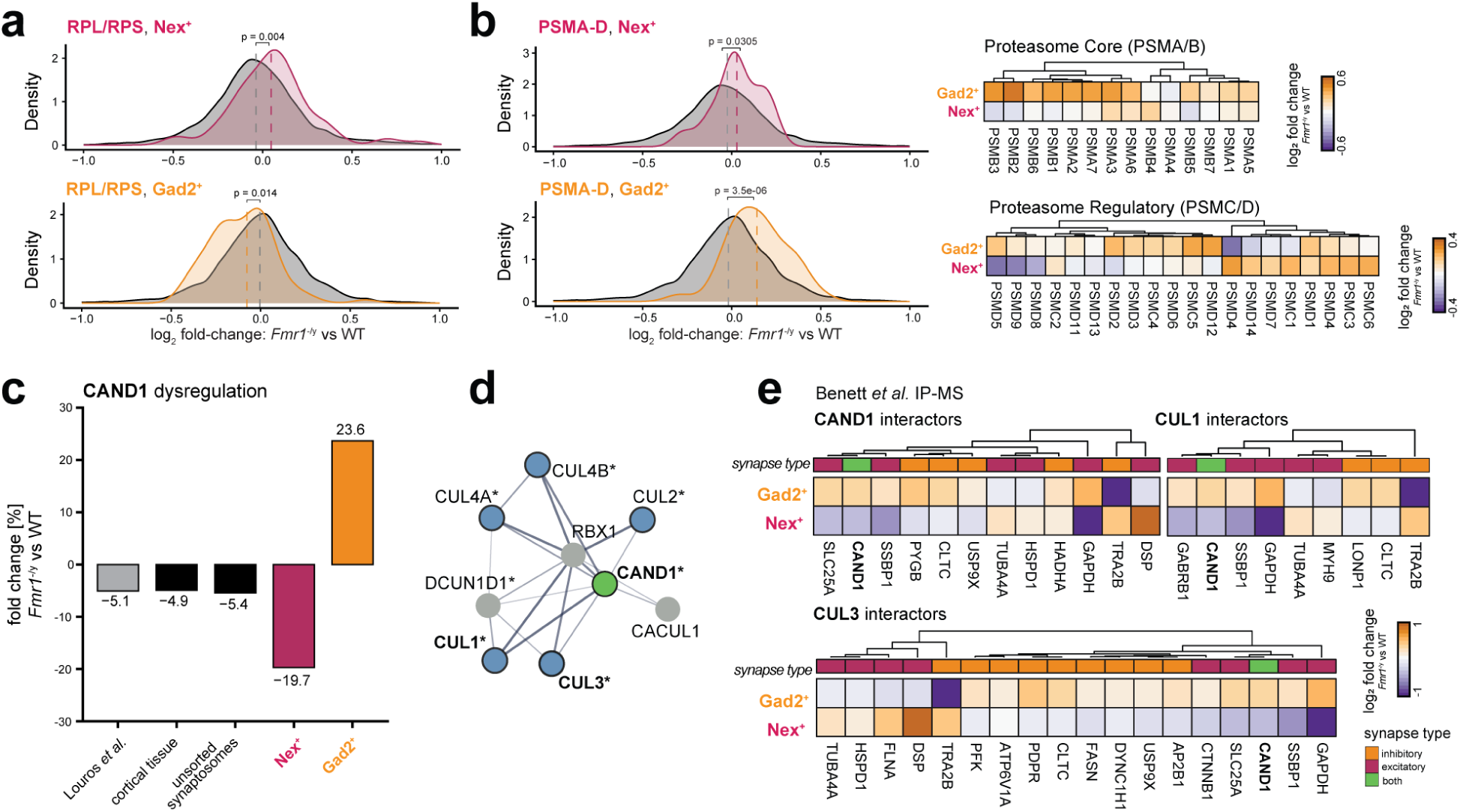
Dysregulation of proteostasis components and the reciprocal protein candidate CAND1, a substrate-recognition exchange factor of Cullin RING-Ligases. **a)** Density plots of log_2_ fold changes (*Fmr1*^-/y^ vs WT) comparing abundances of components of the ribosome (RPL/RPS, highlighted by color) to the overall quantification of the excitatory (maroon) and inhibitory (orange) synaptic proteome (grey; unpaired, two-sided Welch’s t-test). The RP abundance increase in Nex^+^ synaptosomes is in line with reports on *Fmr1*^-/y^ CA1 pyramidal neurons using cell type-specific Trap-Seq [81] and ribosome-GO term enrichment via proteomic profiling of *Fmr1*^-/y^ synapse-enriched P2 fractions [27]. **b)** Density plots of log_2_ fold changes (*Fmr1*^-/y^ vs WT) comparing abundances of components of the proteasome complex (PSMA-D, highlighted by color) to the overall quantification of the synaptic proteome (grey), indicating a significant enrichment for the complex in excitatory (maroon) and inhibitory (orange) *Fmr1*^-/y^ synapses (unpaired, two-sided Welch’s t-test), as previously observed in unsorted, hippocampal *Fmr1*^-/y^ synaptosomes [33]. Heatmaps show quantification of the individual candidates, as summarized in the density plots, separating core (PSMA,B; top) and lid (PSMC,D; bottom) components. **c)** Barplot of significant dysregulation (*Fmr1*^-/y^ vs WT) of the reciprocally-affected candidate CAND1 in excitatory and inhibitory synapses (**Table S16**). Abundance changes in hippocampal *Fmr1*^-/y^ synaptosomes (grey, unsorted; [33]), CX tissue homogenate or unsorted CX synaptosomes (both significant, **Figure S1**, **Table S1**, **Table S2**) are also displayed. **d)** As a modulator of substrate receptors of Cullin-RING ligases (CRLs), CAND1 (green) is reported to interact with Cullin proteins (CULs, blue) as shown in the protein interaction network (StringDB, *Mus musculus*). Interactors quantified in the synapse subtype-proteome (Table S16) are highlighted (*). Edge width reflects the interaction score (StringDB combined score: 0.4-1). **e)** Heatmap of log_2_ fold changes for *Fmr1*-KO (*Fmr1*^-/y^) vs WT of proteins that are significantly dysregulated upon *Fmr1*-KO in either or both synapse types (including relaxed group; Table S16) and also reported CAND1, CUL1, and CUL3 interactors as defined by IP-MS experiments from Benett *et al.* ([69]; IP-MS, 293T cells).

### FMRP loss at excitatory and inhibitory synapses in a sex-specific context

In contrast to hemizygous male individuals with FXS, heterozygous females show variability in reduced FMRP expression and phenotypic severity [70, 28, 31, 29]. We next applied our multi-omic FASS approach to heterozygous *Fmr1^-/+^*mice, mimicking the clinically observed *Fmr1*-genotype of females. Interestingly, *Fmr1^-/+^* mice exhibited a >3-fold reduction of FMRP levels in the cortex and unsorted, cortical synaptosomes, where FMRP levels varied across individual *Fmr1^-/+^* animals similar to clinical observations (**Figures S1a, S6a**). In addition, *Fmr1^-/+^* cortical neurons exhibited a mosaic FMRP expression pattern (**Figure S6b**).

Applying FASS-RNA-Seq to *Fmr1^+/+^* (WT) and *Fmr1^-/+^* mice, we identified ∼7,000 transcripts (**Figure 6a**) in both WT and *Fmr1^-/+^* excitatory and inhibitory synapses as observed for the *Fmr1^-/y^* dataset. PCA revealed a remodelling of the transcriptome (**Figure 6a**) upon reduced FMRP levels for both synapse types. While DEA of the inhibitory synaptic transcriptome yielded a reduced number of dysregulated transcripts compared to *Fmr1^-/y^* mice (∼2,6-fold less: 56 up-, 4 downregulated), excitatory synapses, interestingly, showed a ∼3-fold increase, i.e. 352 up- and 184 downregulated (**Figure 6b, Table S13**). GO analysis (**Figure 6c, Table S20**) of significantly upregulated transcripts in excitatory synapses involved previously annotated FMRP targets and disease-risk genes (**Figure 6d**) revealing an enrichment for synapse organizational, transcriptional and ribosome-associated terms (*Syngap1, Grin2b, Brd4, Kmt2d, Rps28, Rps29, Rpl27*), while downregulated transcripts were particularly enriched for ubiquitin protein ligase binding (*Pink1, Ubxn1, Hspa9*). Inhibitory synapses showed a significant downregulation of transcripts associated with regulation of Ca^2+^-entry, miRNA and nucleobase metabolic processes (*Trpc4, Snd1, Ttr*). More dysregulated FMRP targets and disease-risk genes were found in excitatory synapses (**Figure 6d**). Compared to *Fmr1^-/y^* mice, the relative proportion of dysregulated FMRP targets and disease-risk genes was, however, reduced in both synapse types (6-10% FMRP targets, ∼7% NDD, ∼12% SFARI genes, **Figure 6d, Table S13**). As observed for *Fmr1^-/y^*mice, the transcriptomic dysregulation did not correlate with somatic transcriptomic alterations in both synapse types (R_Nex+_= 0.036, R_Gad2+_ = -0.0032, **Figure S6c**) confirming compartment-specific transcriptomic alterations.

**Figure 6:**
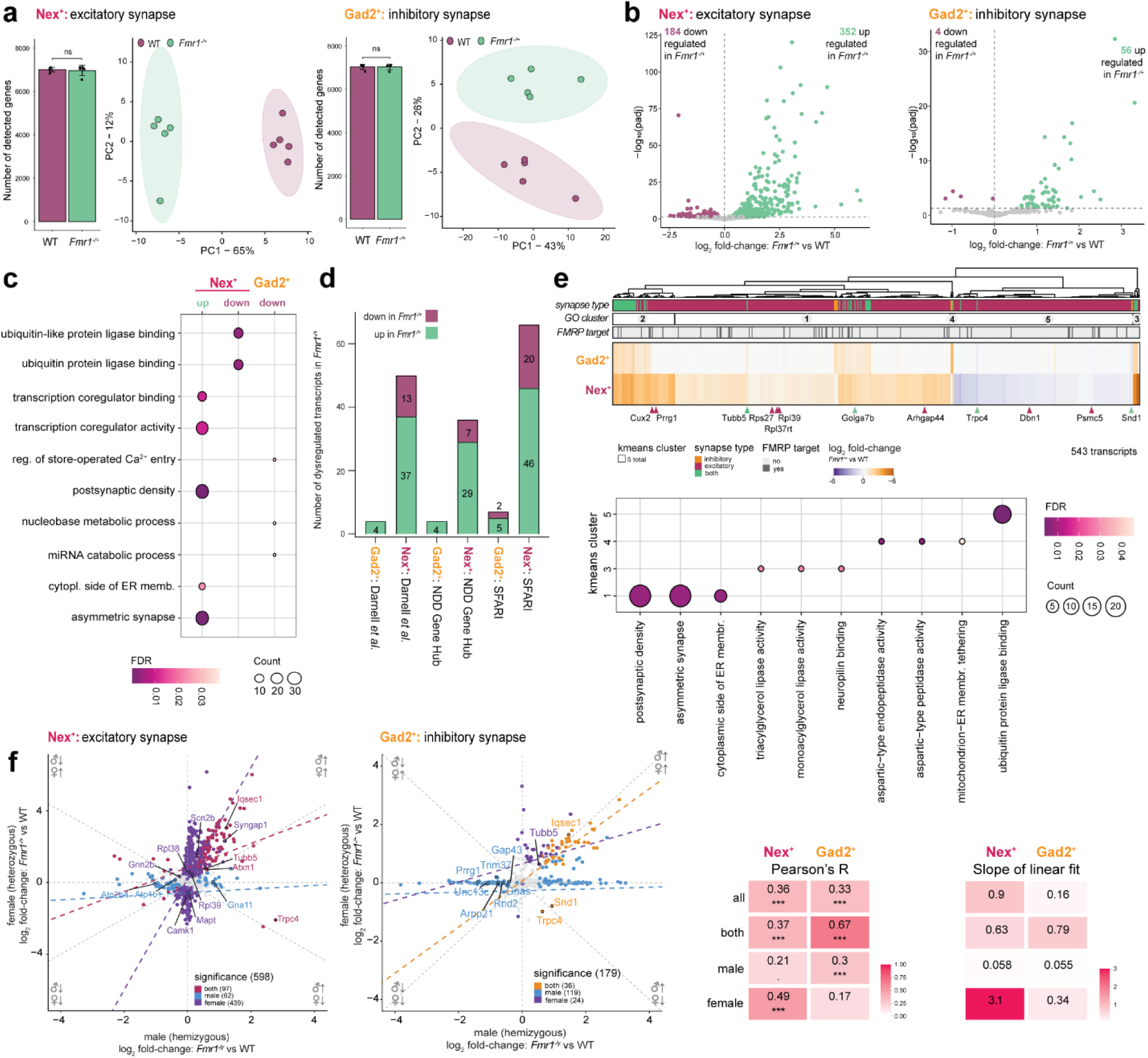
Sex-specific differences and the reciprocal mRNA candidate *Trpc4*, a non-selective cation channel involved in calcium signaling. **a)** Barplots show quantification of ∼7,000 genes in the preparations of both Nex^+^ (left) or Gad2^+^ (right) *Fmr1^+/+^* (WT) and *Fmr1^-/+^* sorted synaptosomes, with no significant difference for the pathological state of the synapse (unpaired, two-sided Welch’s t-test). PCA indicated clear separation of WT and *Fmr1^-/+^* in Nex^+^ and Gad2^+^ sorted synaptosome transcriptomes (top 500 most variable genes). n = 5 animals per condition. **b)** Volcano plots show results of differential transcript (left) abundance analyses for excitatory (top) and inhibitory (bottom) synapses from *Fmr1^-/+^* mice. Both synapse types present with significant abundance changes (FDR < 0.05) of the synaptic transcriptome (Nex^+^: 536, Gad2^+^: 60 transcripts). Dysregulation at the mRNA level appears much more pronounced for excitatory synapses of *Fmr1^-/+^* animals than for inhibitory synapses. **c)** Dotplot shows significant results of GO analyses (FDR < 0.05) of significantly up- and downregulated genes for both synapse types (GO terms from MF, BP, CC). **d)** Number of significantly dysregulated transcripts in *Fmr1*^-/y^ mice annotated as previously reported FMRP mRNA targets [15], NDD-risk genes [53] and ASD-risk genes (SFARI database, [52]) for both synapse types. **e)** Heatmap of log_2_ fold-changes for *Fmr1*^-/+^vs WT of significantly dysregulated transcripts upon *Fmr1*-KO in both synapse types (top). Significant regulation according to synapse type, results of a segmentation analysis (5 clusters), and FMRP mRNA target status [15] are conditionally highlighted below the dendrogram. The results of GO analyses for clusters yielding significant terms (1, 3-5) are summarized in the dot plot (bottom) (top 3 significant terms from GO MF/CC/BP are shown with FDR < 0.05). **f)** Scatterplots of log_2_ fold-changes for *Fmr1*-KO vs WT of significantly dysregulated transcripts upon *Fmr1*-KO in male (*Fmr1*^-/y^, hemizygous vs *Fmr1*^+/y^) and female (*Fmr1*^-/+^, heterozygous vs *Fmr1*^+/+^) animals for both synapse types (Nex^+^ left, Gad2^+^ right). Dysregulated transcripts were highlighted according to significance: significant in both (intersection of male and female) or exclusively in male or female comparisons. The heatmap shows Pearson correlation coefficients and extracted slope-values from linear fit of log_2_ fold-changes for *Fmr1*-KO vs WT of significantly dysregulated transcripts in *Fmr1*^-/y^ and *Fmr1*^-/+^ mice for both synapse types. Significant hits were separated according to their source (union/all, intersection/both, exclusive for male or female).

To understand if the *Fmr1^-/+^* dysregulated transcripts showed a shared or synapse type-specific expression pattern, we performed an unsupervised hierarchical clustering of the dysregulated transcripts followed by GO analyses of the resulting five clusters (**Figure 6e**; **Table S21**). As expected, the two largest clusters showed exclusive up-(cluster 1, 2) or downregulation (cluster 5) of excitatory transcripts confirming the enrichment for synaptic, ribosomal and ubiquitin-associated terms (observed in **Figure 6d**). Interestingly, most upregulated (50/56, cluster 1-4) or downregulated transcripts (3/4, cluster 5) of inhibitory synapses were similarly dysregulated in excitatory synapses. Commonly upregulated transcripts were enriched for lipid metabolism and neuropilin binding (cluster 3: *Abhd2*, *Sema4b*) but also for cytoskeletal organization, vesicle formation and deubiquitination (cluster 1: *Tubb5*; cluster 2: *Iqsec1*, *Otub1*). Transcripts exclusively upregulated in inhibitory synapses formed the last cluster (cluster 4: *Psen2, Catspere2*) while only the transcription factor *Tshz2* showed exclusive downregulation in inhibitory synapses (cluster 5). This indicates that the majority of dysregulation observed in inhibitory synapses was shared with excitatory synapses while the latter showed a more pronounced dysregulation of transcripts essential for synaptic signalling and proteostasis.

Next, we investigated how the hemizygous and heterozygous *Fmr1*-KO genotypes correlated with each other. Integrating the male and female transcriptomic datasets for each synapse type revealed that partial FMRP loss (females) affected excitatory synapses to a greater extent than inhibitory ones. Gad2^+^ synapses showed a 5-fold greater number of transcripts that were exclusively significantly dysregulated in males compared to those in females, indicating a dosage-dependent dysregulation pattern (**Figures 6f, S6d**). In contrast, for Nex^+^ synapses, a small number of transcripts was exclusively dysregulated in males compared to females. For commonly dysregulated transcripts in synapses of males or females (**Figure 6f**), there was high correlation and comparably strong effect size in inhibitory synapses (R = 0.67, slope = 0.79), but lower correlation in excitatory synapses (R = 0.37, slope = 0.63). These observations suggest type-specific differences in susceptibility to reduced FMRP levels, with inhibitory synapses exhibiting a higher sensitivity. We found that, in both synapse types, genotype-specific candidates encoded proteins linked to synaptic organization/signaling, with additional Nex^+^-specific effects on ribosome-associated transcripts, indicating genotype-specific dysregulation of synaptic function (**Figure 6f**). While the majority of commonly dysregulated transcripts was upregulated in both synapse- and genotypes, only the ASD-risk gene *Trpc4* encoding a Ca^2+^-permeable ion channel, showed reciprocal dysregulation across genotypes in both Gad2^+^ and Nex^+^ synapse, i.e. an upregulation in males and a downregulation in females (**Figure 6f**). Overall, we noted that the genotype-specific transcript dysregulation could not be explained by genomic imprinting (**Figure S6e**) or a dysregulation of genes located on sex chromosomes (**Figure S6f**).

Similar to *Fmr1*^-/y^ in males, proteomics analysis of *Fmr1*^-/+^ in females showed only a subtle remodelling of the local proteome of excitatory and inhibitory synapses, with 15 and 21 proteins significantly changed (**Figure S7a-c; Table S16**), respectively. Segregation analysis coupled to GO analyses of the dysregulated subsets revealed perturbation of affected cellular pathways similar to those observed for males (**Figure S7d; Table S22**). Contrasting regulated proteins identified in either males or females directly, we noted that there were 3.6- and 8.2-fold more proteins exclusively dysregulated in males than females for Nex^+^ and Gad2^+^ synaptosomes, respectively (**Figure S7e-g**). Dysregulated proteins in either males or females correlated well in both synapse types (**Figure S7e,f**). However, in line with FMRP dosage dependency, they showed a lower effect size in heterozygous females (**Figure S7e,f**). Proteins significantly dysregulated in both sexes of the distinct synapse types showed high correlation with comparably large effect sizes between hemizygous and heterozygous mutation (**Figure S7f**), suggesting a dosage-independent saturation effect. An integration of the omics-datasets for *Fmr1*^-/+^ synapses revealed a moderate correlation of transcripts and proteins (**Figure S8a**), in line with previous results. Uncoupled mRNA/protein dysregulation (**Figure S8b**) largely corresponded to the molecular phenotype of the *Fmr1*^-/y^ Nex^+^ and Gad2^+^ synapses. Due to the comparably low level of perturbation observed for the female proteome, the overlap of dysregulation in both omics-paradigms was limited to two candidates at excitatory synapses (**Figure S8c**). Inspection of reported FMRP targets showed no uniform dysregulation in either subtype (**Figure S8c**), similar to the observation of the fully FMRP-deficient synapses.

## Discussion

Here, we resolved and integrated local changes in mRNAs and proteins in cortical FMRP-deficient synapses. Our characterization of molecular changes upon *Fmr1*-KO attributed specifically to excitatory or inhibitory synapses provides valuable information in the light of FXS-associated excitatory-inhibitory imbalances in the neuronal circuitry [19, 20].

We characterized the transcriptome and proteome of inhibitory and excitatory WT and *Fmr1^-/y^* synapses, quantifying the abundances of ∼7,000 mRNAs or ∼4,000 proteins for each sample and revealing subtype-specific changes in ∼100s of transcripts and proteins in Nex^+^ or Gad2^+^ synapses, respectively. Overall, there was a small, yet significant, overlap between dysregulated mRNAs/proteins and previously reported FMRP targets [15], in line with previous bulk [34, 60] or cell type-specific omics-experiments [38, 39]. This might be attributed to a non-exhaustive target list with different transcriptomic signatures across brain regions, cell types, and developmental stages, as well as compensation or ripple effects of FMRP loss. Nex^+^ and Gad2^+^ synapses showed equal remodelling in the number of transcripts and proteins that significantly changed upon full FMRP loss, however with only few candidates significantly dysregulated in both synapse types. Notably, most changes in our synapse type-specific analyses deviated from predictions based on abundance values from whole cell or somatic *Fmr1*-KO data.

Which molecules and pathways affected by FMRP loss are similar or unique in different synapse types? Only a minority of mRNAs/proteins showed shared dysregulation at excitatory and inhibitory synapses - the majority was uniquely regulated at one subtype. This general phenomenon of cell type-dependent dysregulation upon FMRP loss concurs with recently reported trends from RiboTag-Seq results of Camk2a^+^ and Pvalb^+^ neurons [39]. Even though the dysregulated mRNAs/proteins differed across synapse types, we noted some degree of pathway-level convergence in our proteomics analysis, i.e. downregulation of similar synapse-associated GO-terms for both Nex^+^ and Gad2^+^ synapses. The transcriptomic data, however, did not point to a comparable, shared GO-enrichment, indicating a greater spread of regulated transcripts across cellular pathways in both synapse types. Overall, even for our relaxed candidates, transcriptomic changes observed at one synapse type showed little to no regulation in the other; proteomic adaptations yielded mostly opposing patterns in excitatory and inhibitory synapses.

Inhibitory *Fmr1*-deficient synapses were predominantly affected by dysregulation of proteins associated with mitochondrial homeostasis, a known central mechanism dysregulated in FXS-models [63–65]. For example, we found opposing dysregulation of the novel protein target SAMM50 (Gad2^+^: up, Nex^+^: down), providing direct connection between a central element of mitochondrial protein synthesis [62] and cristae formation [71] and synapse type-specific changes in an FXS-model. The Nex^+^-exclusive increase of VDAC1 (relaxed candidate) suggests mitochondrial impairment in excitatory, FMRP-deficient synapses, where excessive VDAC1-dependent ER-mitochondria coupling and mitochondrial Ca²⁺ uptake compromise mitochondrial function, as previously demonstrated [64]. Excitatory synapses also showed effects on cellular pathways involved in synaptic function. The different altered mRNA abundances observed in excitatory and inhibitory synapses suggest the potential for type-specific dysregulated local translation and synaptic plasticity (e.g. [25–27]). Generally, local transcript abundance was largely uncoupled from corresponding changes in protein. A few exceptions included uniform upregulation of *Map2*/MAP2 and downregulation of *Aldh7a1*/ALDH7A1 in excitatory synapses. Matched RiboTag-Seq data from Camk2a^+^ neurons concur with our protein-level observation [39], while scRNA-Seq analysis [8] did not show a dysregulation of *Map2,* highlighting the importance of analyzing synaptic compartments. Changes in astrocyte-associated *Aldh7a1*/ALDH7A1 may reflect a global, cell-wide feedback effect of FMRP loss leading to an ALDH7A1-induced decrease in the formation of astrocytic interactions/ tripartite synapses, matching reports on the reduced astrocyte association in tripartite hippocampal synapses in *Fmr1*-deficient mice [23]. Overall, these examples emphasize how FMRP loss specifically dysregulates mRNAs/proteins in inhibitory and excitatory synapses, potentially leading to the impairment of E/I balance.

Which molecules might contribute to the cell type-specific remodelling? We observed a synapse type-independent increase in proteasome subunits together with synapse type-distinct dysregulation of the candidate CAND1. It acts as an exchange factor for CRLs, the largest class of E3 ubiquitin ligases [67], which are associated with various neurological disorders [72, 73]. Trends for CAND1 downregulation were apparent in previous bulk proteomics analyses of *Fmr1*-KO synaptosomes [27, 33]; the small effect sizes in these studies are likely due to averaging opposing regulation. CAND1 dynamically controls the association of CULs with substrate-receptor (SR) complexes of CRLs and is thereby well poised to influence substrate selection, ubiquitination, and ultimately degradation [74, 75, 68, 76]. Beyond CAND1, we noted synaptic localization (without altered abundance upon FMRP loss) of CULs and of other components of the CRL machinery, such as members of the COP9 signalosome or CRL-SRs. Consequently, altered CAND1 abundance would not necessarily result in a uniform in-/decrease in protein degradation alone. Rather, it could contribute to a type-specific remodeling of ubiquitin-dependent proteostasis by altering the composition of SR-repertoire within the CRL complex [74, 75, 68, 77]. This substrate specification in combination with the general maladaptive increase of the protein degradation machinery in *Fmr1*-deficient synapses [33] of both types may therefore lead to type-specific local degradation profiles. Interestingly, candidates from our protein-level analysis (including relaxed group) overlapped with reported interactors of CAND1 and CULs [69] and consistently showed disparate dysregulation in Nex^+^ and Gad2^+^, underscoring CAND1’s potential to fine-tune CRL-complex targeting in a type-specific manner. Furthermore, proteomics studies of bulk *Fmr1^-^*^/y^ synaptosomes blocking proteasomal degradation [33] revealed that e.g. CUL3-interactor CTNNB1 accumulates when proteasomal clearance is inhibited. CTNNB1 is strongly linked to NDDs [78] and to the regulation of (excitatory) postsynaptic strength [79] and the pool of synaptic vesicles [80]. Generally, increased local protein degradation could provide one explanation for unmatched/ upregulated transcripts and proteins (e.g. *Sod1*/SOD1 in excitatory synapses).

How do mRNA and protein adaptations in synapses of heterozygous, female animals compare to those of fully FMRP-deficient males? Related to the “averaging” issue, the interpretation of sex-dependent effects is also complicated: many molecular studies have focused on male *Fmr1*-deficient animals [81, 27, 60, 33], whereas others have not analyzed sexes separately [34, 8, 38, 39]. Findings from hemizygous males should not be extrapolated to heterozygous females, where partial and/or mosaic FMRP expression may produce distinct, non-linear molecular responses across cells and synapses [82, 83]. In our *Fmr1*^-/+^ data, FMRP dosage did not drop linearly (∼3-fold in unsorted synaptosomes) upon KO and was heterogeneous between female mice. Interestingly, with respect to our system-wide investigations, the degree of cell type-specific dysregulation - at Nex^+^ *Fmr1^-/+^* synapses in particular - was much greater at the mRNA than protein level. Our proteomic data suggest a less severe phenotype for the heterozygous *Fmr1*-KO compared to a full FMRP depletion. Few proteins were affected at similar levels in male and female animals. These “threshold-dependent” candidates may regulate pathways that are already maximally perturbed when FMRP is partially lost, e.g. the aforementioned candidate VDAC1. Dosage-dependency was also observed, at least in part, in our transcriptome data, particularly for inhibitory synapses. In contrast, excitatory *Fmr1^-/+^* synapses exhibited a 3-fold increase in the number of dysregulated transcripts compared to *Fmr1^-/y^* mice, affecting mRNAs essential for synaptic signalling and proteostasis. Increased dysregulation of transcripts in *Fmr1^-/+^* compared to *Fmr1^-/y^* mice has previously been reported upon bulk RNA-Seq analyses of the telencephalon of E14.5 mice [84] and hippocampi of 2 month old *Fmr1^-/-^* mice [85]. Why reduced FMRP levels led to an increased dysregulation in excitatory, but not in inhibitory synapses remains to be elucidated. However, we can exclude the impact of genomic imprinting [55] or a predominant dysregulation of sex chromosomal genes. Interestingly, we observed a genotype-specific reciprocal dysregulation of *Trpc4*, a known ASD-risk gene [52], in both synapse types. Trpc4 is activated upon activation of G-protein coupled receptors [86, 87] which leads to a sustained depolarization block, inhibiting neuronal firing [86]. Thus, an increase of *Trpc4*, as observed in synapses of *Fmr1^-/y^* mice, might represent compensation for the hyperexcitability of cortical networks reported in *Fmr1^-/y^* mice [19]. In turn, loss-of-function mutations in *TRPC4* result in developmental delay [88] and homozygous *Trpc4*-KO in male mice results in reduced dendritic arborization associated with social deficits [87]. Thus, a downregulation of *Trpc4*, as observed in *Fmr1^-/+^* mice, could contribute to the clinical FXS symptomatic phenotype.

In summary, we applied our FASS pipeline to a *Fmr1*-KO mouse model to study mRNA and protein remodelling in young adult mice of both sexes and revealed cellular mechanisms underlying the synaptopathy FXS in a compartment- and cell type-resolved manner. We described synaptic dysregulation in response to full and partial FMRP loss that was distinct for excitatory and inhibitory neurons and also not a passive reflection of cell-wide/ somatic effects. Our work (accessible via our interactive platform syndive.org) underscores the importance of synapse type-specific disease research and paves the way for novel treatment strategies in the future.

## Acknowledgements

We thank the MPIBR scientific support facilities: the Animal facility, in particular Emily Northrup, Nina Vogt, Sabahudin Dizdarevic, Silke Zeissler and the rodent team for their support with breeding/ keeping our mice, Georgi Tushev of the computing facility for implementation of syndive.org, and Christine Bohnstaedt of the imaging facility for assistance with image acquisition. Further, we thank Anita Kulak for project support, Stefano Giandomenico for input on neuronal proteostasis, Elena Ciirdaeva for assistance with sequencing and all members of the Schuman lab for helpful discussions. We like to acknowledge that generative AI (GPT-5.6; OpenAI, San Francisco, CA, USA) was used for language editing, optimization of in-house written R-code, or literature assistance. All output and scientific content was conceived, verified, and edited by the authors, who take full responsibility for the manuscript.

## Data availability

All data associated with this manuscript will be made publicly available or provided in supplemental tables upon publication. Raw RNA-sequencing data for synaptosomes have been uploaded to the NCBI Sequence Read Archive [89] and will be released upon publication. The previously published pipeline for RNA-Seq demultiplexing and quantification is available here: https://gitlab.mpcdf.mpg.de/mpibr/schu/synaptosort. All mass spectrometry data associated with this manuscript have been uploaded to the PRIDE repository [90] via the project accession number PXD083908 and will be released once published.

## Supplemental figure legends

**Figure S1:**
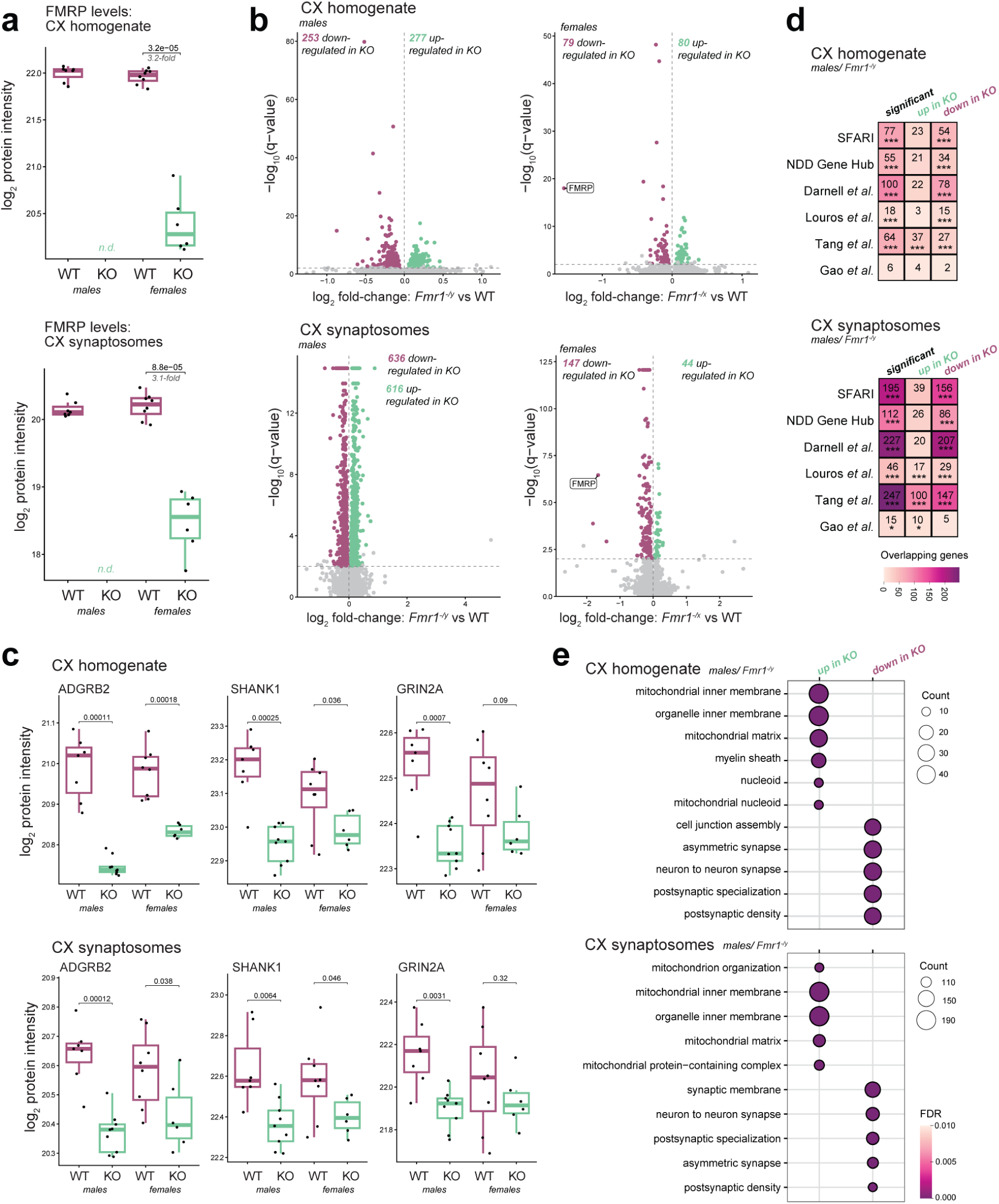
Validation of *Fmr1*-KO (*Fmr1*^-/y^ and *Fmr1*^-/+^) model mice. **a)** Mass spectrometry-based FMRP quantification in *Fmr1*-KO animals, highlighting absence and significant decrease in log_2_ protein intensities of FMRP in cortex homogenate (top) and cortical synaptosomes (F2/3 fraction, bottom), of hemizygous male (*Fmr1*^-/y^) or heterozygous *Fmr1*-KO mice (*Fmr1*^-/+^) and WT (*Fmr1*^+/y^ and *Fmr1*^+/+^) litter mates (unpaired, two-sided Welch’s t-test, p-values listed). n.d. = not detected. **b)** Volcano plots of significantly dysregulated proteins in cortex homogenate (top; **Table S1**) and cortical synaptosomes (bottom; **Table S2**) of hemizygous male (left) and heterozygous female (right) *Fmr1*-KO mice compared to respective WT (downregulation in maroon, upregulation in cyan; FDR < 0.01). **c)** Boxplots show significant decrease of selected, previously reported *Fmr1*-KO affected proteins in cortex homogenate (top) and cortical synaptosomes (bottom), comparing WT (maroon) and *Fmr1*-KO mice of both sexes (unpaired, two-sided Welch’s t-test, p-values listed). **d)** Heatmaps visualize the overlap assessment of all significantly dysregulated, upregulated, or downregulated proteins of hemizygous, male animals as described in b) for cortex homogenate (top) and synaptosomes (bottom) with known ASD-risk genes (SFARI database, [52]), risk genes of neurodevelopmental disorders (NDD Gene Hub, [53]), and previously reported FMRP mRNA targets [15]. Comparisons with previously published proteomics analyses of *Fmr1*-KO mice using unsorted hippocampal synaptosomes (postnatal day 25 - 32, [33]), neocortical synaptosomes (postnatal day 17, [34]), and prefrontal cortex homogenate (3 weeks, [60]) also showed significant overlap with all regulated and particularly downregulated proteins (one-sided Fisher’s exact test; adjusted * p < 0.05, *** p < 0.001). See methods for details. **e)** Dot plot summarizes the results of GO analyses for upregulated (cyan) and downregulated (maroon) proteins from the statistical analyses of cortex homogenate (top; **Table S2**) and synaptosomes (bottom; **Table S4**) from hemizygous male animals as shown in b). The top-5 terms for each protein group are listed (FDR < 0.05). Dot size reflects the number of matched proteins and color indicates significance (FDR).

**Figure S2:**
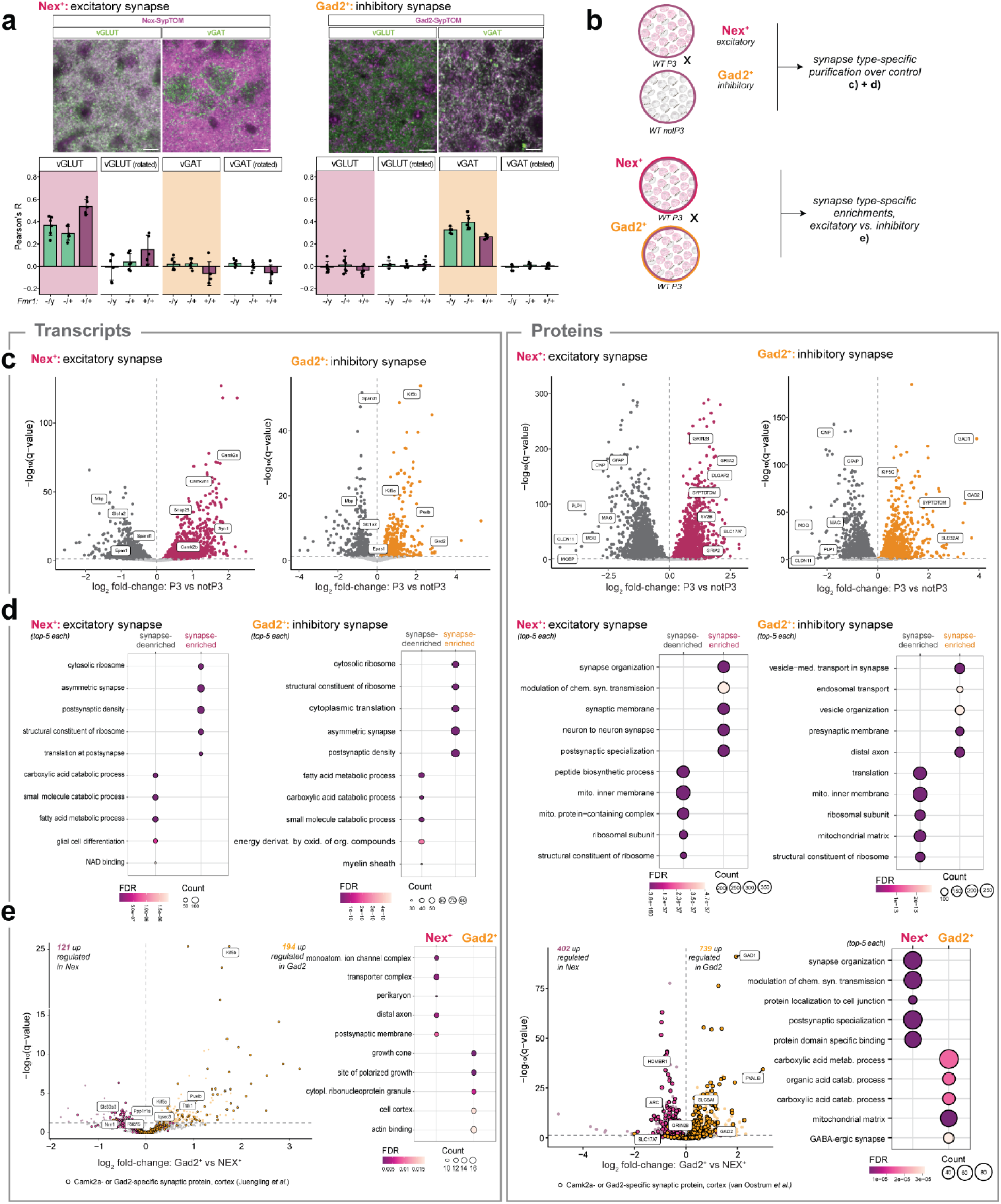
Validation of labelling specificity and purity of fluorescence-activated synaptosome sorting. **a)** Representative images (40x objective) of immunostained cortical brain slices from a Nex-Cre::condSypTOM^+^ (*Fmr*^-+/+^) and a Gad2-Cre::condSypTOM^+^ mouse (*Fmr1*^-/+^), co-staining SypTOM with the excitatory presynaptic marker vGLUT1 and the inhibitory, synaptic marker vGAT to confirm the synapse-type-specific labelling (top). Scale bar = 10 µm. Barplots (bottom) summarize Pearson correlation analyses of the immunostainings of all genotypes, underscoring co-localization of SypTOM with the respective cell type-specific synapse marker for WT, heterozygous *Fmr1*^-/+^ (female), and hemizygous *Fmr1*^-/y^ (male) animals. Analyses of rotated images were used as control. Error bars show standard deviation of six replicate measurements (fields of view). **b)** Analysis strategies to assess synaptic purity (top) based on FASS preparations (P3 FASS gate vs “not-P3” FASS) for both synapse types or cell type-specific enrichments (bottom; Nex^+^ vs Gad2^+^). WT animals were used to define and compare the “undisturbed” synaptic transcriptome and proteome. **c)** Volcano plots comparing the sorted synaptosome fraction (P3 gate) from Nex-Cre::condSypTOM^+^ (maroon) and Gad2-Cre::condSypTOM^+^ (orange) mice with their respective remaining fraction of sorted particles (“notP3”) for mRNAs (left; **Table S5**) and proteins (right; **Table S9**). Enrichment of known excitatory and inhibitory markers (e.g. mRNA: *Camk2a* or *Gad2*; protein: SLC17A7/vGLUT1 or GAD1/2) and depletion of remaining glial ones (e.g. mRNA: *Mbp*, *Sparcl1*; protein: GFAP, CNP) is indicated by gene name label (FDR < 0.05). As expected, protein levels of the SypTOM (used for sorting) are significantly enriched in both preparations. **d)** GO analyses of the synapse-enriched or -deenriched mRNAs or proteins identified in c) are summarized in dot plots. For both transcriptomics (**Table S6**) and proteomics (**Table S10**), the top-5 GO terms (GOCC/MF/BP) ranked by lowest FDR were shown in the plot (FDR < 0.05). Dot size reflects the number of matched transcripts or proteins and color indicates significance (FDR). See methods for details. **e)** Volcano plots of DAAs contrasting excitatory versus inhibitory synaptosomes (P3 gates) to identify the contribution of synapse type-enriched transcripts and proteins in the WT synaptic transcriptomes (**Table S7**) and proteomes (**Table S11**). No systematic pattern for synapse type-preferential dysregulation was found in our disease-associated findings attributing the synapse-type specific changes observed in *Fmr1*-KO mice to FMRP deficiency rather than the synapse type of origin. Significant differences in quantity of excitatory (e.g. mRNA: *Nrn1*, *Ppp1r1a*, *Rab15*; protein: ARC, SLC17A7/ vGLUT1) or inhibitory (e.g. mRNA: *Kif5b*, *Pvalb*; protein: SLC6A1/ vGAT, GAD1/2, PVALB) candidates are highlighted by color (FDR < 0.05). For mRNAs, overlap with previously reported excitatory and inhibitory synaptic transcriptome [1] is highlighted in the volcano plot by a black outline. For proteins, overlap with the synapse-type-enriched proteins of the previously reported excitatory and inhibitory synaptosome proteome [3] is also indicated by a black outline. GO analyses of the Nex^+^- or Gad2^+^-enriched mRNAs (**Table S8**) or proteins (**Table S12**) are summarized in dot plots (left to respective volcano plot) for top-5 GO terms of significantly enriched GOCC/MF/BP terms (FDR < 0.05). Dot size reflects the number of matched proteins and color indicates significance (FDR).

**Figure S3:**
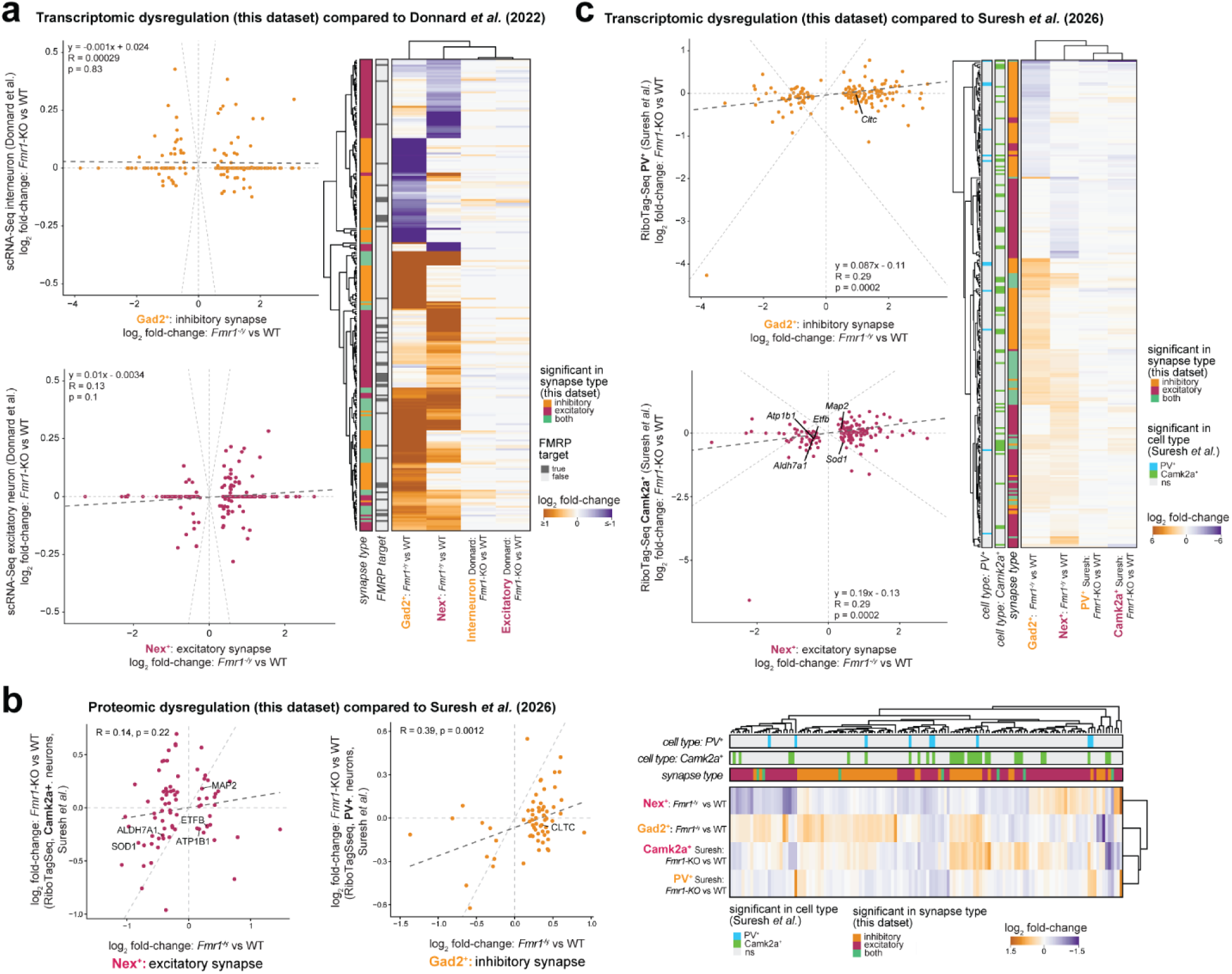
Single-cell RNA-Seq and RiboTag-Seq datasets of *Fmr1*-KO mice show poor to modest correlation with synaptic *Fmr1^-/y^* datasets. **a)** Heatmap shows a combined clustering of log_2_ fold-changes of *Fmr1*^-/y^ vs WT of significantly dysregulated transcripts in Nex^+^ and Gad2^+^ synaptosomes from this study (**Table S13**) compared to regulated somatic transcripts of cortical excitatory and interneurons from *Fmr1*-KO mice defined upon scRNA-Seq analysis (postnatal day 5, male and female animals combined; [8]), contrasting both cell-types and highlighting previously reported FMRP mRNA targets [15]. Scatterplots of the respective log_2_ fold-changes show low correlation between both integrated datasets. **b)** Scatterplots of log_2_ fold changes for *Fmr1*^-/y^ vs WT of significantly dysregulated proteins in Nex^+^ and Gad2^+^ synaptosomes from this study (as defined in Figure 3d**, Table S16**) compared to regulated ribosome-associated transcripts of Camk2a^+^ and Pvalb^+^ neurons from somatosensory and primary visual cortex, as quantified via cell type-specific RiboTag RNA-Seq (postnatal day 70 - 100, male and female animals combined; [39]). Highlighted candidates are selected based on commonly dysregulated mRNA-protein pairs (as introduced in Figure 4c). The heatmap shows a combined clustering of respective log_2_ fold changes, contrasting both cell-types, highlighting known products of FMRP-bound mRNAs [15] and significance of the dysregulation in the work of Suresh and colleagues [39]. **c)** Heatmap shows a combined clustering of log_2_ fold-changes of *Fmr1*^-/y^ vs WT of significantly dysregulated transcripts in Nex^+^ and Gad2^+^ synaptosomes from this study compared to regulated ribosome-associated transcripts of Camk2a^+^ and Pvalb^+^ neurons from somatosensory and primary visual cortex, as quantified via cell type-specific RiboTag RNA-Seq (postnatal day 70 - 100, male and female animals combined; [39]), contrasting both cell-types and highlighting previously identified FMRP mRNA targets [15] and significance of the dysregulation in the work of Suresh *et al.* [39]. Scatterplots of the respective log_2_ fold-changes show low correlation between both integrated datasets. Labelled transcripts, again, indicate commonly dysregulated mRNA-protein pairs.

**Figure S4:**
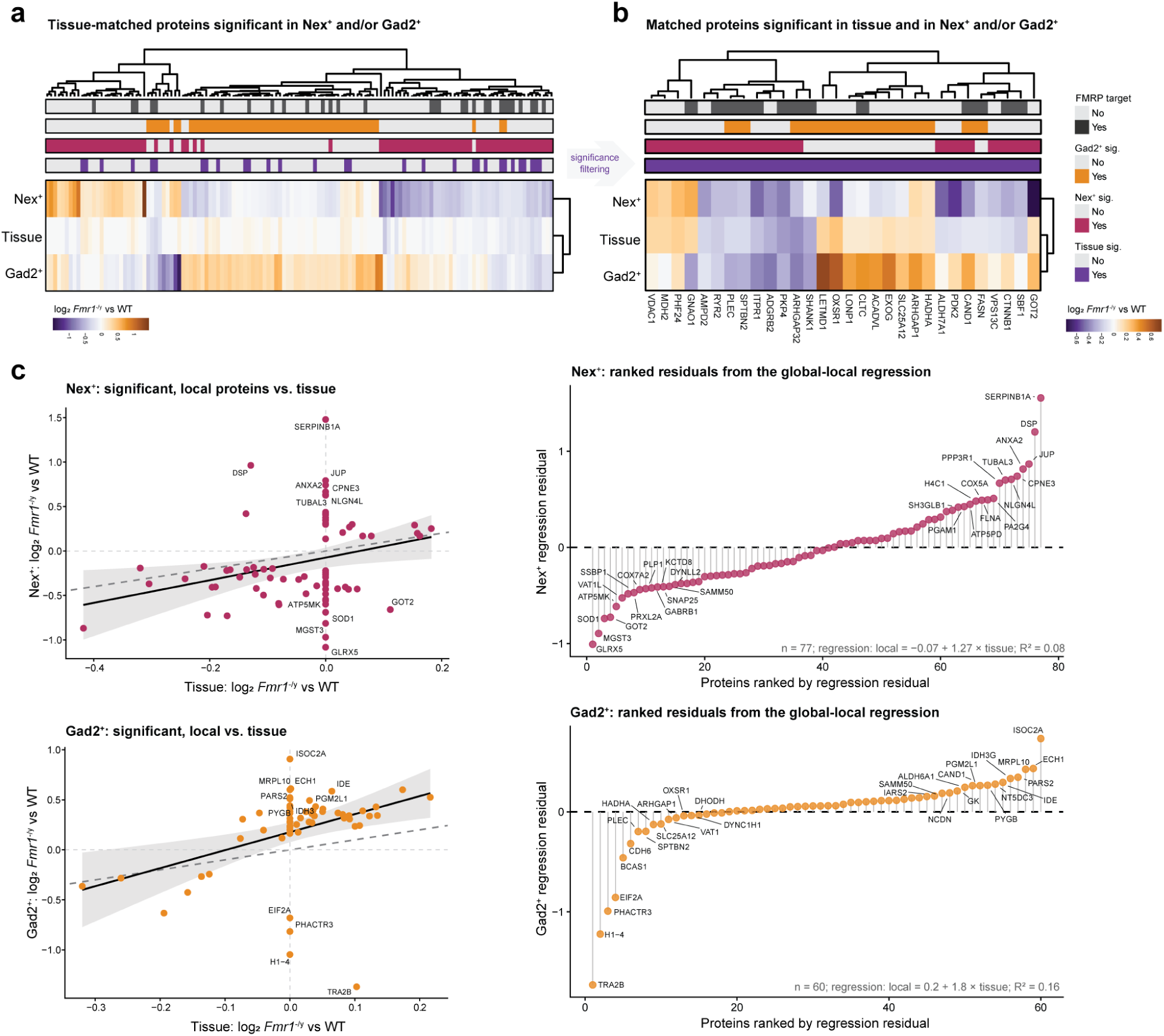
Relationship between protein abundance changes following *Fmr1*-KO in cortex tissue (global) and cell type-specific, cortical synaptosomes (local). **a)** Heatmap (left) summarizes the log_2_ fold changes of proteins significantly dysregulated at the synapse subtype-specific local compartment (excitatory or inhibitory synaptosomes, *Fmr1*^-/y^; as defined in Figure 3d**, Table S16**) to their matching values in cortical tissue homogenate (as introduced in **Figure S1; Table S1**). All proteins that were significantly dysregulated upon *Fmr1*-KO in either synapse-type and had valid quantification at tissue level are shown (**Table S17**). Heatmap (right) filters for significance both globally and locally. Significance is highlighted by color below the dendrogram, as well as the information of them being encoded by FMRP-bound transcripts [15]. **b)** Scatterplots contrast the log_2_ fold changes of proteins significantly dysregulated at the synapse type-specific local compartment to their matched values from cortex homogenate. Linear models were fitted for Nex^+^- (top) and Gad2^+^-significant proteins (bottom) and regression lines with 95% confidence intervals were displayed. Regression residuals were calculated as the difference between the observed and fitted local log₂ fold change, ranked according to amplitude, and thus used as a quantitative measure of protein-specific deviation from the global trend, as shown in the lollipop plots (left).

**Figure S5:**
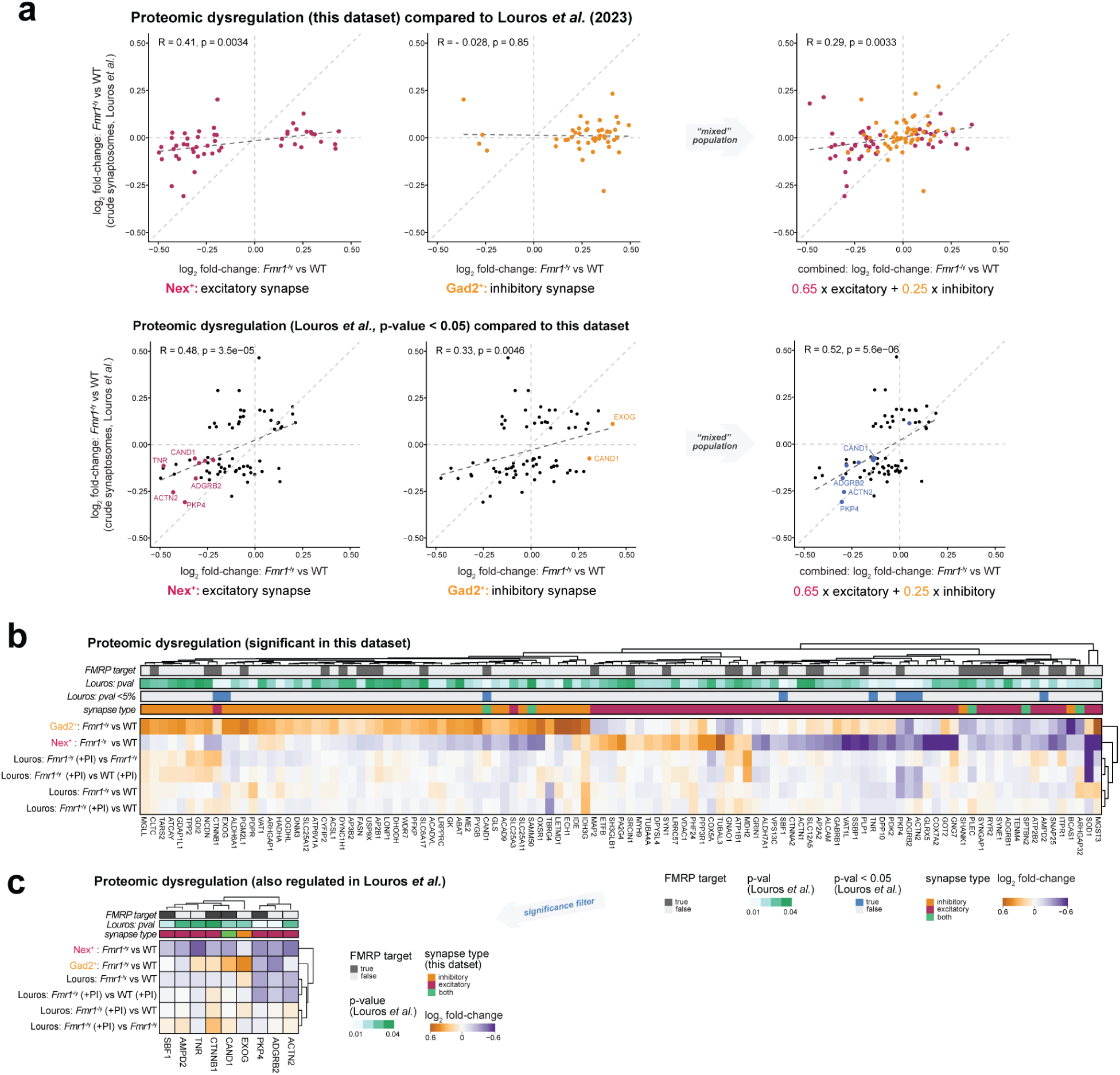
Correlation of synaptic protein dysregulation due to *Fmr1^-/y^* with published data on (cell type-unspecific) synaptic remodelling. **a)** Scatterplots (top) of log_2_ fold changes for *Fmr1*^-/y^ vs WT of significantly dysregulated proteins in this study (as defined in Figure 3d**, Table S16**) compared to protein regulation observed in studies of unsorted hippocampal synaptosomes of *Fmr1*^-/y^ and WT mice (postnatal day 25 - 32; [33]). Cell type-specific regulation upon *Fmr1*-KO (Nex^+^ left, Gad2^+^ middle) and an approximated mixture of proteins from both synapse types reflecting unsorted synaptosomes (right, “mixed population” see Figure 1b for respective ratios) was contrasted. Scatterplots (bottom) present log_2_ fold changes for *Fmr1*^-/y^ vs WT of significantly dysregulated proteins in this study compared to proteins regulated proteins (p < 0.05) in the study of Louros and colleagues [33]. Candidates showing significant regulation in our cell type-specific analyses yielded significant Pearson correlation for excitatory and the “combined” correlation, while there was no correlation for the inhibitory candidates, suggesting a dilution of this regulation in bulk dataset. For candidates regulated in the published work, there is significant Pearson correlation compared to our quantifications across all synapse-types, where the correlation coefficient of excitatory synapses surpasses the one of inhibitory synapses. As expected, the combination approximating a mixed population (see Figure 1b) shows the highest correlation to the data by Louros and colleagues. Poorer correlation of our synapse-type specific protein changes to the previously published data than vice versa and the enhancement upon assuming a “mixed population” are suggestive of a masking/dilution effect in the bulk synaptosome analysis. **b)** Heatmaps of log_2_ fold changes of significantly dysregulated proteins in this study (relaxed group) compared to all proteins quantified by Louros and colleagues or **c)** all regulated proteins (p < 0.05; *Fmr1*^-/y^ vs WT) from unsorted synaptosomes [33], including log_2_ fold changes from additional analyses of unsorted synaptosomes generated from hippocampal slices treated with (+PI) or without the proteasome inhibitor MG132. As reported in the study, proteins showing an accumulation in *Fmr1*^-/y^ synapses without proper proteasomal clearance may indicate their dependence on accelerated proteasomal degradation upon FMRP loss, such as CTNNB1.

**Figure S6:**
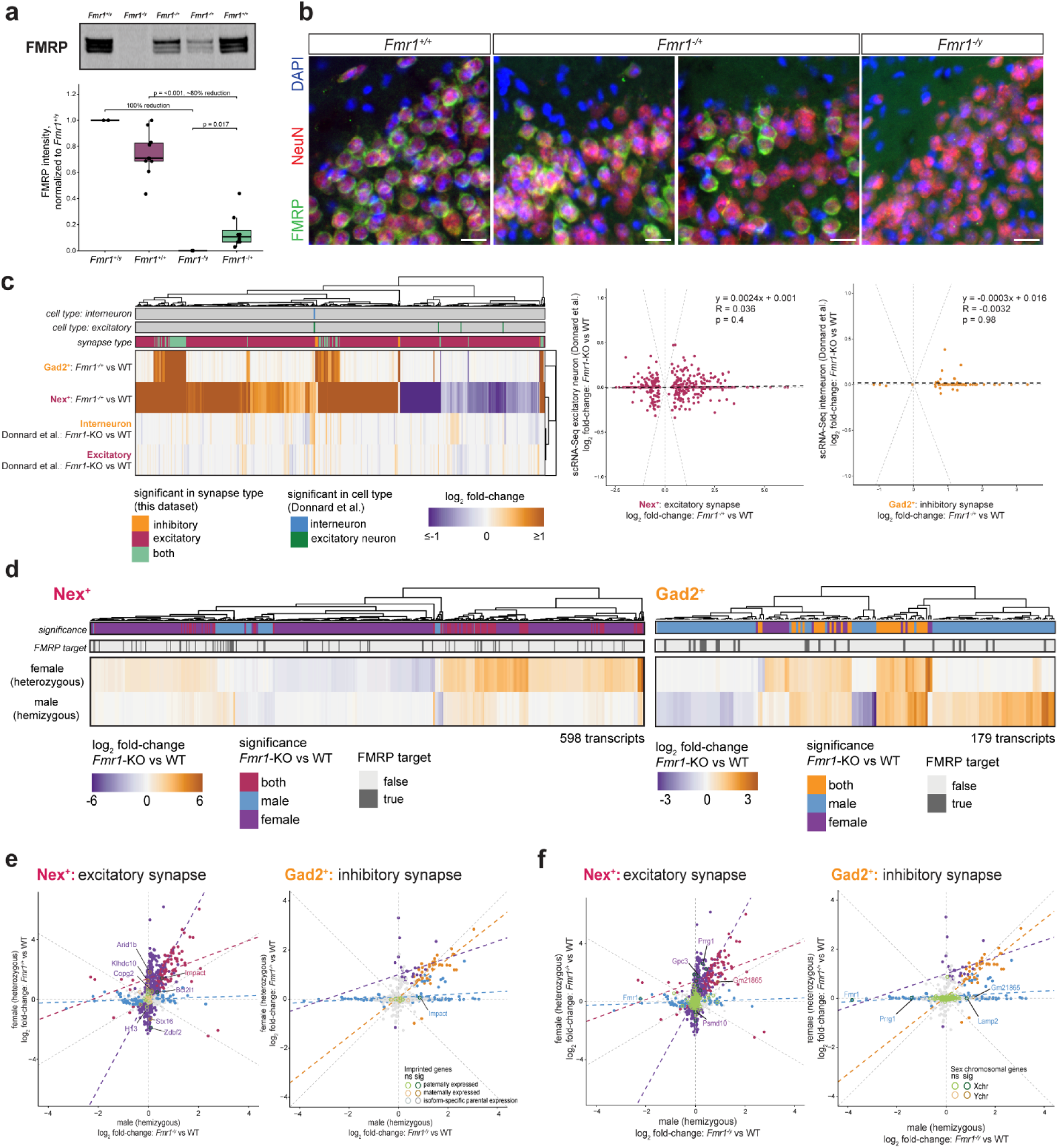
Transcriptomic profiling of subtype-specific synapses in heterozygous (*Fmr1*^-/+^), female animals. **a)** Western blots of *Fmr1^+/y^* (n = 1 animal, 2 technical replicates), *Fmr1^-/y^* (n = 1 animal, 2 technical replicates), *Fmr1^+/+^* (n = 10 animals) and *Fmr1^-/+^* (n = 8 animals) cortex homogenates showed full depletion of FMRP in *Fmr1^-/y^*. In *Fmr1^-/+^* mice, FMRP levels varied across individual animals. For quantification, FMRP-levels of *Fmr1^-/y^*, *Fmr1^-/+^,* and *Fmr1^+/+^* were normalized to *Fmr1^+/y^*. Statistical significance was assessed using an unpaired, two-sided Welch’s t-test. **b)** Representative maximum intensity projected images (40x objective) of immunostainings for FMRP (green) and NeuN (red) in cortical brain slices (border between cortical layer 1 and layer 2) of *Fmr1^+/+^* (n = 1), *Fmr1^-/+^* (n = 2) and *Fmr1^-/y^* (n = 1) animals showed full depletion of FMRP in neurons in *Fmr1^-/y^* mice whereas *Fmr1^-/+^*animals showed a mosaic FMRP expression pattern in neurons. Scale bar = 20 µm. **c)** Heatmap of significantly dysregulated transcripts in Nex^+^ and Gad2^+^ synaptosomes of *Fmr1^-/+^* mice annotated with scRNA-Seq data of cortical excitatory and interneurons of *Fmr1*-KO mice [8] (left). Scatterplots show dysregulation of significantly dysregulated transcripts of excitatory synapses (Nex^+^) and somata [8] (right) and of inhibitory synapses (Gad2^+^) and somata [8] indicating low correlation between subcellular compartments. **d)** Heatmap summarizing the regulation described in Figure 6e while highlighting FMRP mRNA target status [15]. **e)** Male-female scatter plot with annotated imprinted genes [55]. The male-female discrepancy observed in each synapse type cannot be explained by the dysregulation of imprinted genes. Imprinted genes that are not significantly dysregulated in the *Fmr1*-KO transcriptomic datasets are highlighted in light green for paternally expressed, in light yellow for maternally expressed and in light grey for isoform-specific paternal expression of imprinted genes. Significantly dysregulated imprinted genes are shown in the respective dark color. **f)** Male-female scatter plot with annotated sex chromosomal genes for each synapse type. The male-female discrepancy observed in each synapse type could not be explained by the dysregulation of sex chromosomal genes. X-chromosomal genes (Xchr) that were significantly dysregulated are highlighted in dark green and Ychr genes in dark yellow. Not significantly dysregulated sex chromosomal genes are shown in the respective light color.

**Figure S7:**
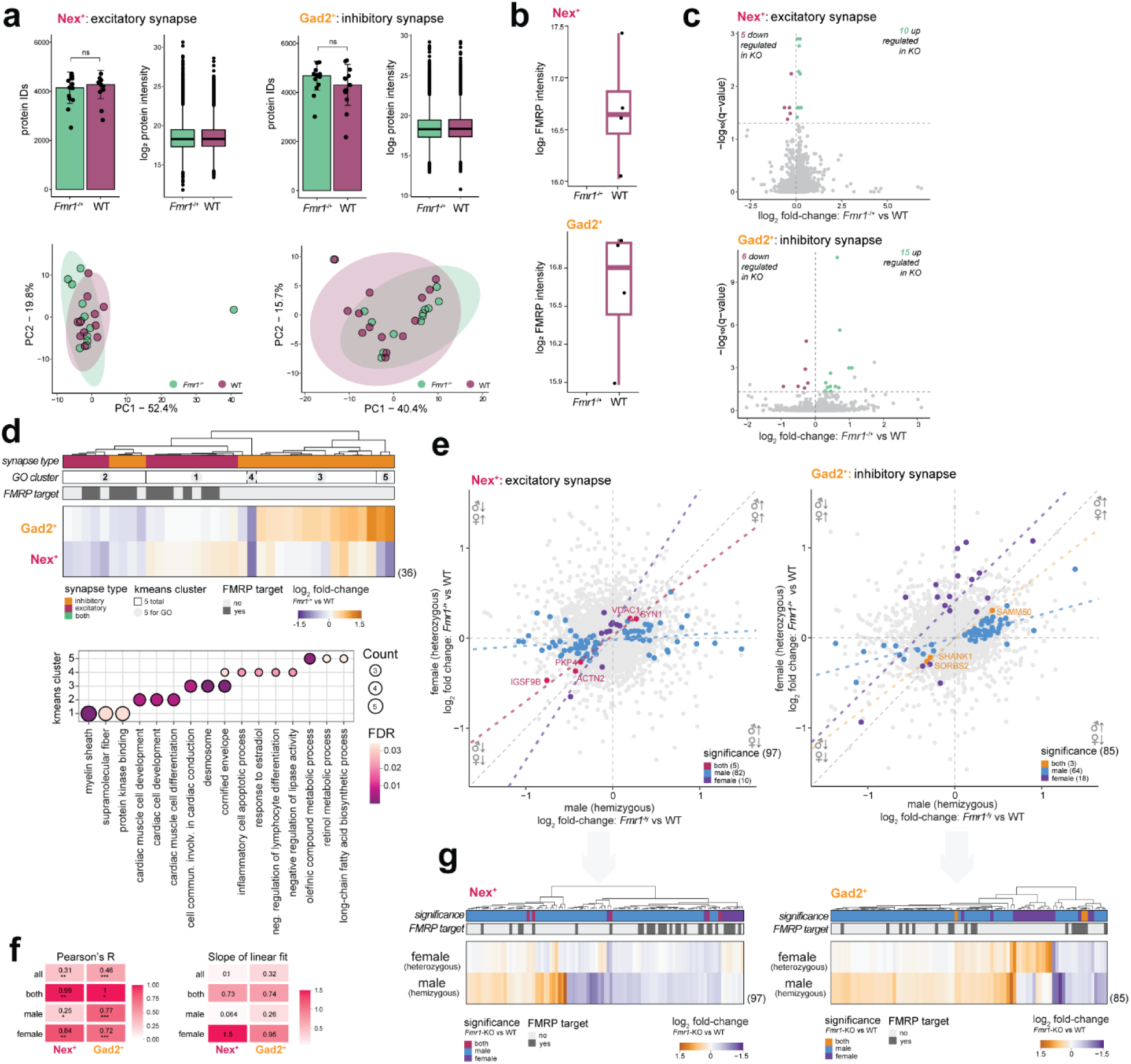
Proteomic profiling of subtype-specific synapses of *Fmr1*^-/+^ female mice. **a)** Barplots show quantification of ∼4,000 proteins in the preparations of both Nex^+^ (left) or Gad2^+^ (right) sorted synaptosomes, with no significant difference for the pathological state of the synapse of heterozygous females (unpaired, two-sided Welch’s t-test). High similarity is also observed for boxplots of overall protein intensities in both synapse types and conditions. PCA indicates no clear separation of WT (*Fmr1*^+/+^) and *Fmr1*-KO (*Fmr1*^-/+^) in Nex^+^ or Gad2^+^ sorted synaptosome proteomes (top 500 most variable proteins, log_2_ protein intensity). **b)** Boxplots of FMRP levels (log_2_ protein intensity) show its abundance at each synaptic compartment of female WT mice and its absence in synapses of KO mice. **c)** Volcano plots show results of differential protein abundance analyses for excitatory (top) and inhibitory (bottom) synapses from *Fmr1^-/+^* mice (**Table S16**). Both synapse types present with significant abundance changes (FDR < 0.05) of its local proteome (Nex^+^: 15, Gad2^+^: 21 proteins) upon FMRP loss. **d)** Heatmap of log_2_ fold changes for *Fmr1*-KO vs WT of significantly dysregulated proteins upon *Fmr1*-KO in female animals and both synapse types (top). Significant regulation of synapse type, results of a segmentation analysis (kmeans clustering, n = 5), and FMRP mRNA target status [15] are conditionally highlighted below the dendrogram. The results of GO analysis for clusters yielding significant terms are summarized in the dot plot (bottom, FDR < 0.05; **Table S22**). **e)** Scatterplots of log_2_ fold changes for *Fmr1*-KO vs WT of significantly dysregulated proteins upon *Fmr1*-KO in male (*Fmr1*^-/y^, hemizygous vs *Fmr1*^+/y^) and female (*Fmr1*^-/+^, heterozygous vs *Fmr1*^+/+^) animals for both synapse types (Nex^+^ left, Gad2^+^ right). Dysregulated proteins were highlighted according to significance: significant in both (intersection of male and female) or exclusively in male or female comparisons. **f)** Heatmaps of Pearson correlation coefficients (left) and extracted slope-values (right) from linear fits of log_2_ fold changes for *Fmr1*-KO vs WT of significantly dysregulated proteins upon *Fmr1*-KO in male (hemizygous) and female (heterozygous) animals for both synapse types (Nex^+^ left, Gad2^+^ right). Again, significant hits were separated according to the significant comparison (union/all, intersection/both, exclusive for male or female). **g)** Heatmap summarizing the regulation described in e) while highlighting previously reported FMRP mRNA target status [15].

**Figure S8:**
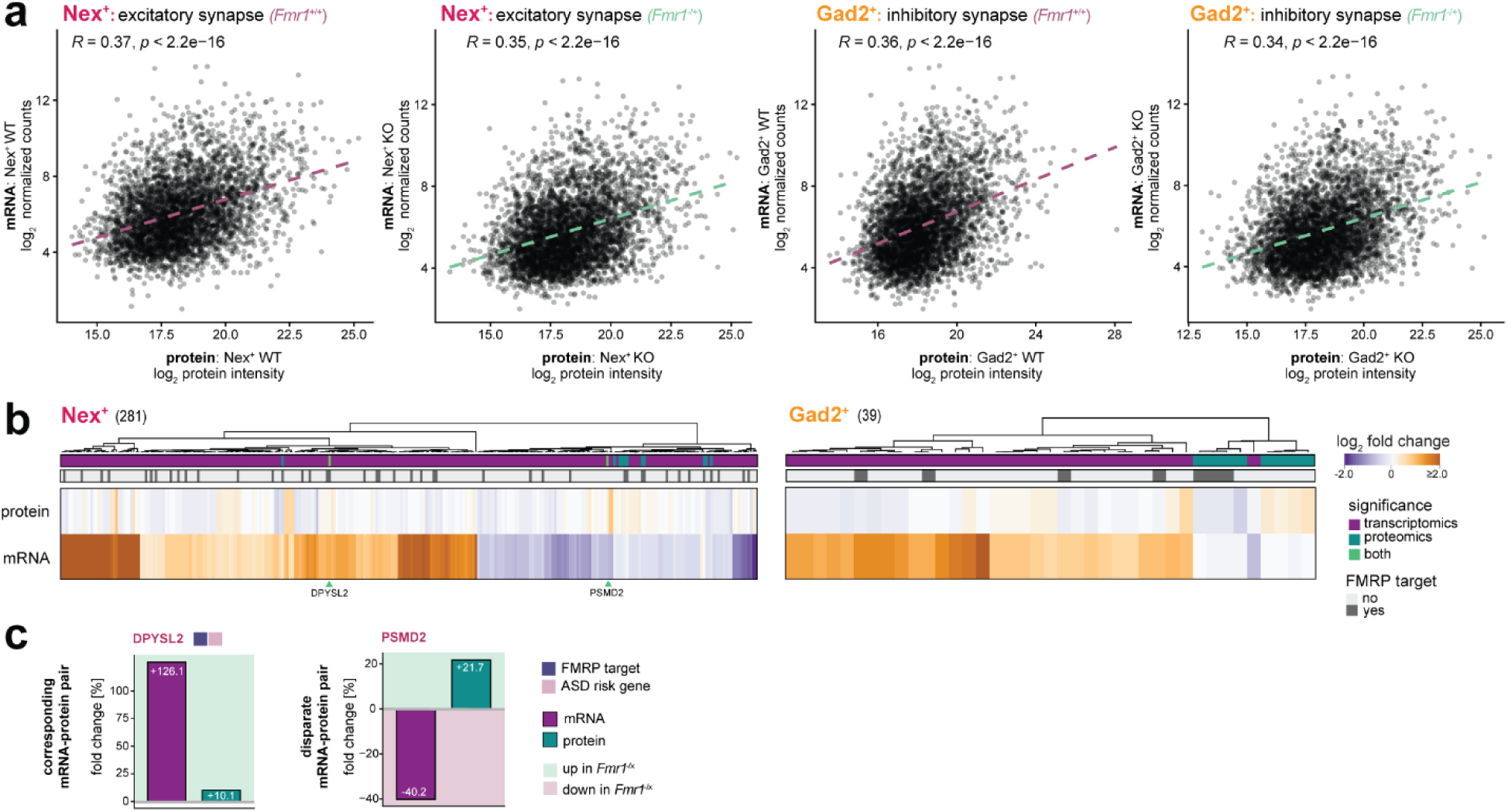
Differences in *Fmr1*-KO (*Fmr1*^-/+^) associated mRNA and protein dysregulation at excitatory and inhibitory synapses of female animals. **a)** Scatterplots show correlation of mean abundances of mRNAs (log_2_ DESeq normalized counts) and their respective protein (log_2_ protein intensity) for WT (*Fmr1*^+/+^) and *Fmr1*-KO (*Fmr1*^-/+^) animals of excitatory (left) and inhibitory (right) synapses, quantified by Pearson’s correlation and highlighted via linear fit. **b)** Heatmaps show log_2_ fold changes (*Fmr1*-KO vs WT) of significantly dysregulated mRNAs and proteins. Significance of dysregulation at transcript and/or protein level and reported FMRP mRNA target status [15] are conditionally highlighted below the dendrogram. Candidates significantly dysregulated in both omics-regimes are labelled. For visualization, the log₂ fold change scale was restricted to the indicated range, accounting for differences in effect-size magnitude between the datasets. **c)** Barplots of the shared mRNA-protein pairs show corresponding and disparate regulation using the fold change (in percent). Color of the candidate name indicates dysregulation observed in Nex^+^ or Gad2^+^ synaptosomes. Functional annotation with respect to previously reported FMRP targets [15] or ASD-risk genes [52] is provided next to the gene name.

